# Population genetic structure and range limits of *Prostanthera cineolifera* (Lamiaceae), a vulnerable shrub with a patchy distribution

**DOI:** 10.1101/2023.03.27.534300

**Authors:** Ruth L. Palsson, Ian R.H. Telford, Jeremy J. Bruhl, Rose L. Andrew

## Abstract

Integrating molecular data is essential for clarifying the distributions and genetic structures of species that have histories of misidentification and misapplication of names. There has been confusion about the species limits of the Vulnerable *Prostanthera cineolifera* with respect to morphologically similar specimens in the Hunter Valley, New South Wales Australia and morphologically dissimilar specimens in the Lower Hawkesbury Valley New South Wales and from northeastern New South Wales. To test the species limits of *P. cineolifera* and related taxa specimens were collected from across the range and augmented with herbarium specimens. We used morphometric analysis of 18 morphological characters across 51 operational taxonomic units. Using the DArTseq reduced representation sequencing platform, 9,559 single-nucleotide polymorphisms (SNPs) across 122 individuals were recovered for molecular analysis. Both morphological and molecular analyses produced three concordant clusters (1) *P. cineolifera*, (2) a group sharing similarities with *P*. sp. Hawkesbury (B.J.Conn 2591), and (3) a group allied with *P. lanceolata* and *P. ovalifolia*. These results indicate that the specimens form northeastern New South Wales are more likely to be *P. lanceolata*, not *P. cineolifera*, and that specimens from the Lower Hawkesbury are of an undescribed species with the phrase name *P*. sp. Hawkesbury (B.J.Conn 2591). Within *P. cineolifera* there was pronounced genetic differentiation among populations. Little evidence of inbreeding was observed, but the newly recognised, more isolated populations had the lowest genetic diversity. This study provides new information about the range of the species and its genetic structure that informs the conservation priorities for this species.

## Introduction

A primary goal of conservation is to maintain the evolutionary potential of species (Frankel 1974; Bowen 1999; Milot *et al*. 2020). This goal can be hampered by a lack of information on the distribution and genetic diversity of species (e.g. Gitzendanner *et al*. 2012; Neel and Ellstrand 2003; Coates *et al*. 2018). Taxonomic uncertainty can impede conservation efforts, in two ways. First, undiscovered and undescribed species are easily overlooked in conservation planning and decision-making (Dubois 2003). Second, misidentification of populations can make species appear more widespread than they are or can risk misallocating conservation efforts (Mason *et al*. 2016; Conn *et al*. 2021; Collins *et al*. 2022). In addition to characterizing genetic diversity and connectivity, population genetics can be used to clarify the boundaries between species and test the identity of questionable populations (Ottewell *et al*. 2016)

Disjunct species distributions raise many questions about the biology and conservation of species (Llorens *et al*. 2015; Ottewell *et al*. 2016; Millar and Byrne 2020), stimulating research on the cause of the disjunction and potential for local adaptation. However, disjunct populations may have simply been misidentified, possibly as a result of unsuccessful identification keys or a lack of robust taxonomic work on the group (examples in Williams *et al*. 2006; Sassone *et al*. 2021; Wilde and Barrett 2021). The question of the taxonomic identity of the disjunct populations must be asked: are they the same species or different?

This issue is particularly acute for populations that appear to be at the edge of the range of a species. Small, range-edge populations may be at risk of losing genetic diversity by genetic drift and inbreeding and therefore have an increased extinction risk (Frankham *et al*. 2012). Edge of range populations could also be of high conservation value as they may contain unique genetic diversity (Razgour *et al*. 2013) that may play an important role in adaptation of the species to environmental change such as climate change (Kyrkjeeide *et al*. 2020). However, we risk over-investing or under-investing in such populations if species limits are poorly understood.

Developments in sequencing technologies have accelerated population genetics research in the conservation context (Hunter *et al*. 2018). Reduced representation sequencing approaches can generate robust single nucleotide polymorphism (SNP) datasets without the need for a time-consuming development stage (Andrews *et al*. 2016). Analysis of reduced representation sequence data does not rely on the availability of a reference genome, making the data uniquely suitable for studying the population structure, genetic diversity and species boundaries of non-model organisms (da Fonseca *et al*. 2016; Rossetto *et al*. 2021), as well as testing the identity of questionable populations or unknown samples (Renner *et al*. 2022).

*Prostanthera* Labill. is an Australian endemic genus of shrubs (a few species growing to low trees). The genus includes 114 described species and 18 accepted phrase names (Australian Plant Census IBIS database 1970; Wilson *et al*. 2019; Conn *et al*. 2021; O’Donnell *et al*. 2021) and are mostly shrubs, some of which are so-called species complexes with morphological variability and wide distributions such as the *Prostanthera lasianthos* Labill. species complex (Conn 1993) and *P. ovalifolia* R.Br. *s.l*. (Althofer 1978). An integrative taxonomic approach has been successful in partially resolving the *P. lasianthos* species complex (Conn *et al*. 2021), but confusion remains within the *P. ovalifolia* complex. For example, *P. cineolifera* R.T.Baker & H.G.Sm. has been frequently confused with *P. ovalifolia* and *P. lanceolata* Domin. Although it is listed as a “vulnerable” species under both Commonwealth (Environment Protection and Biodiversity Conservation Act 1999) and New South Wales (Biodiversity Conservation Act 2016) legislation, the genetics and species limits of *P. cineolifera* have not yet been examined.

The species limits of *P. cineolifera* need resolution to inform effective management of the species (Office of Environment & Heritage n.d.). There has been confusion about the species limits of the Vulnerable *Prostanthera cineolifera* with respect to morphologically similar specimens in the Hunter Valley, New South Wales and morphologically dissimilar specimens in the Lower Hawkesbury Valley New South Wales and from northeastern New South Wales. The distribution described in the Approved Conservation Advice (Department of the Environment 2008) masks uncertainty about numerous collections that have been determined variously as *P. cineolifera*, *P. lanceolata* or *P. ovalifolia* (Fig. 1). Morphologically dissimilar specimens have been determined as *P. cineolifera*, while some morphologically similar specimens have been determined as *P. cineolifera*, *P. ovalifolia* or even *P. discolor* R.T.Baker. For example, preliminary examination suggests that those from the Lower Hawkesbury catchment are morphologically dissimilar to those from the Hunter region. Two populations from northeastern New South Wales (Tabbimoble Creek and Pillar Valley) approximately 450 km from the Hunter Valley have also been identified as *P. cineolifera* (AVH 2017) although they occur within the range of *P. lanceolata* (Althofer 1978). If these populations are indeed *P. cineolifera*, they may warrant extra protection because of the likelihood that they possess unique genetic variation or experience distinct threatening processes (e.g. hybridization).

**Fig. 1.**
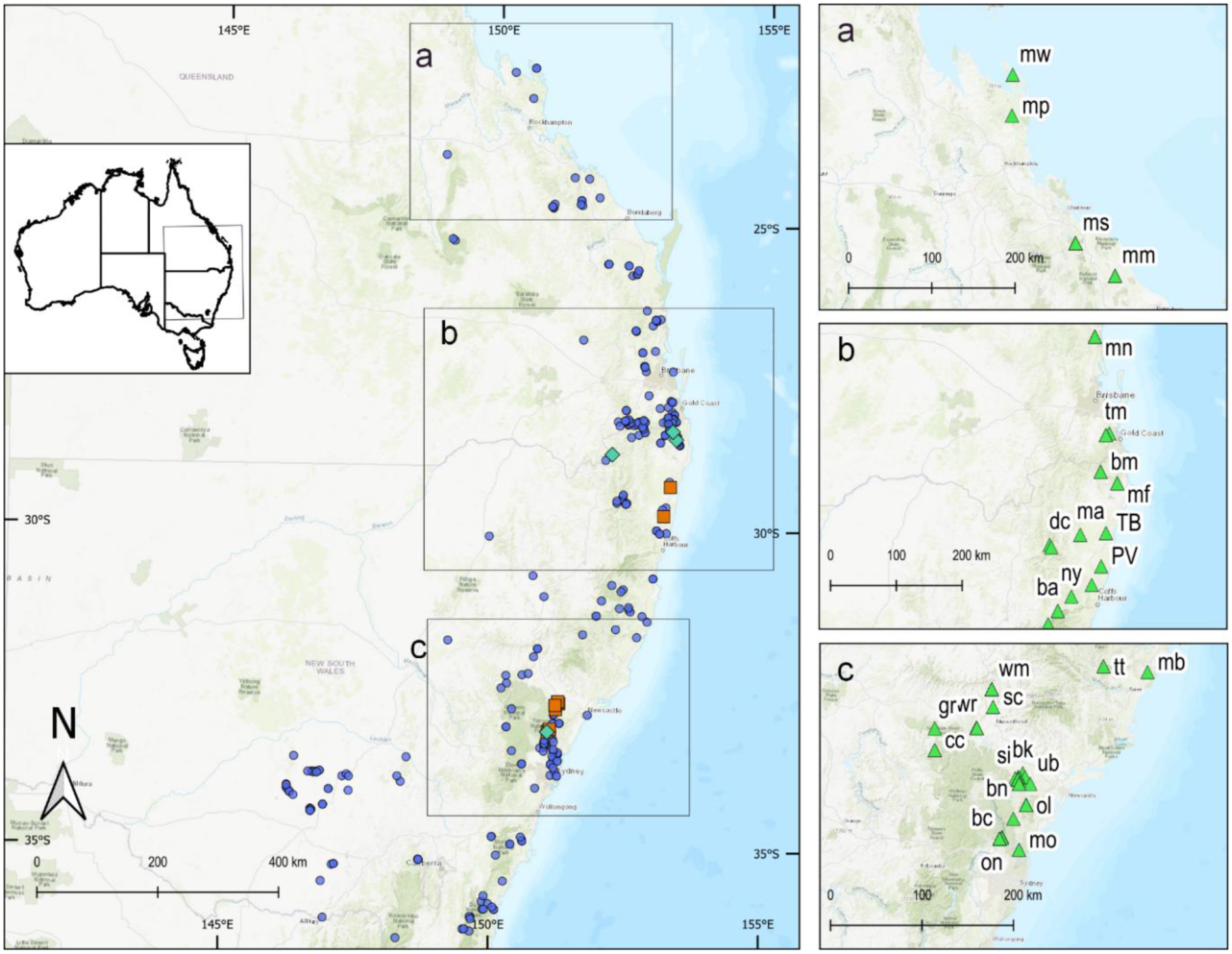
Distribution in eastern Australia of *Prostanthera cineolifera* (orange squares), *P. lanceolata* (aqua diamonds) and *P. ovalifolia* (blue circles) according to the Australasian Virtual Herbarium (AVH; 13 July 2017). Insets of collections used for phenetic and /or molecular analysis: a Central coastal Queensland near the type location of *P. ovalifolia,* b Southeastern Queensland and northeastern New South Wales near the type location of *P. lanceolata* and c the Hunter Valley and Lower Hawkesbury catchment New South Wales near the type location of *P. cineolifera*. (For location codes see Table S1). Note: AVH data were cleaned of cultivated specimens and specimens with geocoding errors that placed them offshore

The distribution of *P. cineolifera* is highly patchy, with populations typically occurring on shallow or skeletal soils, typically on or near sandstone/conglomerate ridges or in gullies towards the western limits of its range. While development and agricultural intensification have occurred in the regions it inhabits, it is unclear how much it has contributed to the patchiness of populations. The focal group of *Prostanthera* in this study have entomophilous flowers with the beetle+fly floral type of Wilson *et al*. (2017). These flowers are protandrous, suggesting that outcrossing is favoured (Wilson *et al*. 2017). Populations of *Prostanthera* vary from scattered individuals (Palsson 2020) to multi-aged stands to large stands of seemingly a single age cohort that are possibly the result of mass germination after a disturbance such as fire or drought (Althofer 1978 and pers. obs). We thus expect patterns of genetic diversity to vary among populations.

Using field observations, herbarium searches, morphological study and next-generation sequencing-based molecular analysis, we examined variation in *Prostanthera cineolifera* and relatives to address three principal questions. 1) Are the disjunct populations in northeastern New South Wales (Tabbimoble Creek and Pillar Valley) and in the Lower Hawkesbury River region isolated, ecologically distinct populations of *P. cineolifera*, or do they belong to different, perhaps undescribed species? This question has bearing on whether urgent protection is needed to preserve unique genetic diversity within the species or new, undescribed species with restricted ranges. 2) To what extent are populations subdivided within *P. cineolifera*, and is it likely to be due to recent fragmentation or long-term isolation due to habitat patchiness? 3) Is there cause for concern about the health of any *P. cineolifera* populations, or the species as a whole? While more work is needed, we also lay the groundwork for further taxonomic assessment of related species by clarifying their species limits with regard to *P. cineolifera*.

## Materials and methods

### Sampling

We used AVH (2017) data for *P. cineolifera*, *P. lanceolata* and *P. ovalifolia* and specimens held at N.C.W. Beadle Herbarium to identify sampling locations to target the New South Wales and southeastern Queensland populations of the study group (Fig. 1). Populations that we determined to be *P. cineolifera* were sampled from 10 sites in four board locations in the Hunter Valley: Wingen Maid Nature Reserve (one), Wallaby Rocks near Sandy Hollow (one), Cousins Creek in Wollemi National Park (one) and near the type locality in the Broke-Pokolbin area near Singleton (seven). The populations of the two putative outlying collections of *P. cineolifera* from the north coast of New South Wales (AVH 2017) were sampled. We aimed to test whether these populations were *P. cineolifera* or not—perhaps *P. lanceolata*. Other populations of *P. lanceolata* in southeastern Queensland (type location) and northeastern New South Wales were sampled. Populations of *Prostanthera* in the Lower Hawkesbury catchment that have sometimes being determined as *P. cineolifera* were also sampled (Fig. 1 and Table S1). We aimed to include samples from type localities or as close to type localities as possible; however, the type locality for *P. ovalifolia* of Mt Westall, inside the Shoalwater Bay Training Area, was inaccessible. For *P. ovalifolia*, we restricted our sampling to populations close to the type locality of Mt Westall, i.e. Mt Stanley (225 km SSE of Mt Westall) and Mt Maria (270 km SSE of Mt Westall), and to locations in the Hunter and Lower Hawkesbury Valleys where populations of *Prostanthera* had been determined as *P. ovalifolia*.

We refer to the morphologically similar specimens in the Lower Hawkesbury catchment that have been determined to be *P. cineolifera*, or *P. lanceolata* or *P. ovalifolia* as *P.* sp. Hawkesbury, as they are morphologically similar to *P*. sp. Hawkesbury (B.J.Conn 2591) NSW Herbarium. The specimens from Olney State Forest that had been determined to be *P. ovalifolia* can be separated from *P. ovalifolia* and *P.* sp. Hawkesbury on the basis of leaf shape, leaf texture and scent were given the working name *P.* sp. Olney State Forest (R.L. Palsson 166) and are referred to here as *P.* sp. Olney State Forest. The specimens from northeastern New South Wales are referred to by their locations as their species identities were unknown. The final study group included *P. cineolifera, P. lanceolata, P. ovalifolia, P.* sp. Hawkesbury and *P*. sp. Olney State Forest, the populations from Tabbimoble Creek and Pillar Valley plus specimens with the following phrase names (Chapman 2007) which were included because of similarity to one of the previously mentioned entities: *P.* sp. Mt Marsh (L.M.Copeland 3770) and *P*. sp. Oxley Wild Rivers National Park (J.B. Williams NE 91044), hereinafter referred to as *P.* sp. Mt Marsh and *P.* sp. Oxley Wild Rivers National Park respectively.

Fieldwork, under New South Wales Office of Environment and Heritage Scientific Licence SL100305, involved collection of herbarium vouchers and metadata including, soil descriptions, digital images, leaf material for phytochemical and morphological analyses, material to be freeze dried, material stored in silica gel (Si Gel), material for cuttings, and occasionally seedling transplants. All voucher specimens were lodged at NE, with duplicates allocated to other Australian herbaria as appropriate. Population size estimates and the presence of juvenile and adult plants were also recorded. Further field work was undertaken in 2019 and 2022 to search for previously unknown populations of *P. cineolifera.* Field collections were assigned to species based on diagnostic morphological features (Baker and Smith 1912; Domin 1928; Conn 1993), such as stem indumentum, bud and calyx shape. However, the putative *P. cineolifera* populations at Tabbimoble Creek and Pillar Valley were considered ‘unknown’, despite exhibiting similar morphology to *P. lanceolata*.

Morphological analysis included representative specimens from our field collections and from the following herbaria: Queensland Herbarium (BRI), Australian National Herbarium (CANB), N.C.W. Beadle Herbarium (NE), and National Herbarium of New South Wales (NSW). Herbarium codes follow the Index Herbariorum (see http://sweetgum.nybg.org/ih/, accessed 17 July 2022). Herbarium searches revealed specimens morphologically consistent with *P. cineolifera*—all determined as *P. ovalifolia*—from Goulburn River National Park, Manilla, Tamborine Mountains, Queensland (BRI AQ0336457) (see Supplementary Information for details) and Upper Moore Creek (Tamworth) These specimens had the diagnostic features of *P. cineolifera* (features of stem indumentum, bud, and calyx shape) and were included in the phenetic analysis.

### Morphological analysis of *P. cineolifera* species boundaries

Characters used were guided by character lists in Williams *et al*. (2006) and Conn *et al*. (2013), augmented by characters following our examination of specimens. Characters included quantitative (10), unordered multistate (6) and binary characters (2). Quantitative characters were scored as a mean of three measurements. Measurements were made either by steel rule or eyepiece graticule at 5X magnification, calibrated against a steel rule. Initially, information for 18 morphological characters (Table S2) across 51 operational taxonomic units (OTUs) were recorded into DELTA (Dallwitz 1980), then exported via Excel to PATN v4 (Belbin 2004–2013) for phenetic analysis (Table S3). OTUs with more than two missing data points were also removed, leaving 48 OTUs (Isb1-T, Omp1 and Omw1-T, with had 4, 5 and 7 missing data points, respectively).

Morphometric analysis used the Gower metric for association. The flexible unweighted pair-group method with arithmetic mean (UPGMA) with the default β = - 0.1 (Belbin 2013) was used for clustering, and semi-strong hybrid multidimensional scaling (SSH-MDS) was used for ordination with the maximum number of iterations set to 100 with other parameters at their default settings. The Kruskal-Wallis (KW) statistic was used to evaluate the ability of variables to discriminate between a set of groups in the cluster; higher KW values have more discriminating power (Belbin 2004–2018). Principal-component correlation (PCC) values were used to evaluate the contributions of specific variables to the ordination. These values, expressed as *r*^2^, are an indication of the amount of variance accounted for by that character (Belbin 2004–2018).

### Molecular analysis of *P. cineolifera* species boundaries and population structure

For DNA extraction, leaf samples were freeze-dried, dried on Silica gel, or sampled from herbarium specimens (Table S4). Genotyping-by-sequencing using the DArTseq platform was conducted by Diversity Arrays Technology P/L (Canberra, Australia), producing single nucleotide polymorphism (SNP) data. One hundred and fifty-seven samples from Clade J (Wilson *et al*. 2012), including 133 individuals from the study group, were randomized across two plates prior to sequencing (Meirmans 2015). Eighteen of the samples had a technical duplicate in the second plate (see Technical duplicates in Supplementary Information). The resulting 75 bp reads were assembled de novo and variants scored by DArT using their proprietary pipeline.

The data returned from DArT were imported into the statistical software R 4.0.2 (R Core Team 2020) and processed with the package *dartR* v1.9.9.1 (Gruber and Georges 2018). Following Gruber *et al*. (2018), sites were filtered based on repeatability (100% required) and call rate (minimum 95%). After filtering individuals based on individual call rate (minimum 90%), loci with similar trimmed sequence tags (threshold 25%) were removed, resulting in 12,123 loci scored for 148 individuals (for details see Tables S6 and S7 in Supplementary Information). Preliminary ordination of this dataset via principal coordinate analysis (PCA) (Gower 1971) was conducted using the ‘gl.pcoa’ function in *dartR* (Gruber *et al*. 2017).

Scrutiny of the resultant plot indicated four anomalous specimens, likely through errors in plate setup, considering their plate positions. These four specimens were removed along with the technical duplicates in plate 2, leaving 12,107 loci scored for 144 individuals (see Anomalous specimens in Supplementary Information).

To examine the relationships of the populations in the study group, a neighbour-joining tree was constructed among populations across the corrected set of taxa using the command, ‘gl.tree.nj’, in *dartR*, and rooted with Launceston specimens as the outgroup (Gruber *et al*. 2017). Subsequent analyses were limited to either the focal geographic region (9,559 loci scored across 122 individuals), in order to test the species boundaries of *P. cineolifera* in the relevant geographic context, or to samples considered to be *P. cineolifera* based on morphological analyses. Filters (Table S6) were applied as in the larger dataset, except that threshold of the call rate filter on individuals was lowered to 80% to keep as many individuals as possible (Table S7). An additional filter of minor allele frequencies using a threshold of 0.05 was applied as recommended in Frichot and Francois (n.d.) resulting in 6,450 loci in 53 individuals for the within-species dataset. Ordination of the molecular data via PCA was conducted as described above, and non-hierarchical clustering and ancestry estimation were implemented as a complementary approach.

As isolation by distance may drive population structure, it was important to distinguish genetic discontinuities between species from apparent clustering due to patchy sampling across a continuum (Frantz *et al*. 2009). We used the R package ConStruct v1.0.4 (Bradburd *et al*. 2018) for non-hierarchical clustering of populations within a spatially explicit population-level model that estimates ancestry proportions while accounting for isolation by distance. It characterizes the contributing ancestral populations as ‘layers’ in which allele frequencies may vary according to an estimated degree of spatial autocorrelation. Evaluation of model fit via cross-entropy allows the spatial model to be compared with one in which allele frequencies do not vary geographically, as well as comparing different values of *K*, the number of ancestral populations. We included all sampled populations within the study group (*P. cineolifera, P. lanceolata, P. ovalifolia, P.* sp. Hawkesbury and *P*. sp. Olney State Forest, *P.* sp. Mt Marsh, *P.* sp. Oxley Wild Rivers National Park and the populations at Tabbimoble Creek and Pillar Valley that had been determined as *P. cineolifera*).

Individual ancestry coefficients were estimated using sparse nonnegative matrix factorization (sNMF) in the LEA package v3.2.0 (Frichot and Francois 2014). This package uses cross-entropy values to suggest the probable number (*K)* of genetic clusters/ancestral populations in the data and assign individuals to genetic clusters (Frichot and Francois n.d.). The ‘snmf’ command was run with *K* = 1–10 and at least 10 replicates at each value of *K*.

Basic genetic summary statistics were calculated for the putative species in the study region (Table S8) and for populations within *P. cineolifera* using the ‘gl.basic.stats’ command in *dartR* (Gruber *et al*. 2017). Population sample size (*N*), observed heterozygosity and unbiased expected heterozygosity were computed for both datasets, excluding populations with *N* < 5 for within-species analyses. Inbreeding coefficients were not calculated for the species-level analysis, as small population sample sizes in some species meant that hierarchical analyses could not be used to account for population subdivision within species.

Divergence among putative species and among populations (excluding those with *N* < 5) within *P. cineolifera* was compared using pairwise *F_ST_*, calculated with the ‘stamppFst’ command in the *StAMPP* package with 1000 permutations (Pembleton *et al*. 2013). *F_ST_* values less than 0.05 are considered low, those between 0.05 and 0.25 are considered moderate, and those greater than 0.25 are considered pronounced (Freeland *et al*. 2011).

## Results

### Field observations

Only populations within the Hunter Valley region were consistent with morphology of *P. cineolifera*. The morphological boundary between *P. lanceolata* and *P. ovalifolia* was indistinct in the field, so OTU codes were tentative, although they were supported by observations of flowering time. In this group, there is a gradient of flowering time with a discontinuity at the Brisbane River. Thus, unless otherwise stated, *P. ovalifolia* refers to specimens of the *P. ovalifolia/lanceolata* group collected for this study north of the Brisbane River, Queensland, and *P. lanceolata* is used for specimens of the *P. ovalifolia/lanceolata* group collected for this study south of the Brisbane River.

The populations from the Lower Hawkesbury catchment were not consistent with the morphology of *P. cineolifera* but were consistent with the morphology of *P.* sp. Hawkesbury and this name has been used throughout this paper. The population from Olney State Forest, morphologically similar to *P.* sp. Hawkesbury but can be separated in the field by leaf shape, leaf texture and scent, have been referred to as *P*. sp. Olney State Forest throughout this paper. Two populations of *P.* sp. Oxley Wild Rivers National Park were included because of morphological similarity to *P.* sp. Hawkesbury. Following additional field work in 2019 and 2022, a further visit to the National Herbarium of New South Wales and communication with SAJ Bell and KH Stokes in 2021, a total of 22 *P. cineolifera* populations are known. In unexplored parts of Broken Back Range, Goulburn River National Park and Wollemi National Park, there is considerable suitable habitat of slopes, ridge crests and gullies for more potential populations of *P. cineolifera*.

The number of individuals in populations of *P. cineolifera* ranges from three (Broke Road) to c. 1,000 individuals at Bees Nest Ridge 4 and Yellow Rock Rd 4 (Table 1). The estimated minimum population size of *P. cineolifera* was approximately 5,200, as of October 2022 (Table 1). We found no immature plants of *P. cineolifera* on Wallaby Rocks or Broke Road.

**Table 1.** Estimated population sizes and age structure of *Prostanthera cineolifera* populations known to the authors. Multi aged populations generally included a range of ages from young shrubs less than 1 m high to mature shrubs up to 6–7 m; some mature populations lacked the very old shrubs. Shaded populations were not included in any analyses. NP = National Park, NR = Nature Reserve, Rd = Road

| Locality | Collection site | Population estimate | Age structure |
| --- | --- | --- | --- |
| Broken Back Range | Bees Nest Ridge 1 | 30 | most shrubs less than 2 m high |
|  | Bees Nest Ridge 2 | 80 | most shrubs less than 2 m high |
|  | Bees Nest Ridge 3 | 320 | multi-aged |
|  | Bees Nest Ridge 4 | 1,000 | multi-aged |
|  | Broken Back Trail | 100 | multi-aged, but lacking very old shrubs |
|  | Sawpit Rd | 300 | multi-aged, but lacking very old shrubs |
|  | Off Broken Back Rd<sup>a</sup> | 40 | multi-aged |
|  | Yellow Rock Rd 1 | 500 | multi-aged |
|  | Yellow Rock Rd 2 | 45 | most shrubs less than 2 m high |
|  | Yellow Rock Rd 3 | 130 | most shrubs less than 2 m high |
|  | Yellow Rock Rd 4 | 1000 | multi-aged |
|  | Yellow Rock Rd 5 | 400 | multi-aged |
| Broke Cessnock area | Broke Rd<sup>b</sup> | 3 | mature plants only |
|  | Singleton Military Area (two sites) <sup>c</sup> | 100 | multi-aged |
| Goulburn River NP | Morrison's Flat Trail | 500 | multi-aged |
| Scone | Owens Gap<sup>d</sup> | 140 | unknown |
| Sandy Hollow | Wallaby Rocks | 10 | mature plants only |
| Upper Bellbird | Limestone Creek (two sites) <sup>e</sup> | 40 | multi-aged |
| Wingen Maid NR | Wingen Maid | 200 | multi-aged |
| Wollemi NP | Cousins Creek | 300 | multi-aged |
| **Estimated minimum total population** |  | **5,200** |  |

a KH Stokes, *pers. comm.* 2021, b These plants did not survive the severe drought that ended early 2020, c SAJ Bell, *pers. comm.* 2021,
d NSW 785391, e LMC Foster, *pers. comm.* 2018

All other known populations of *P. cineolifera* were multi-aged including immature plants. Some populations (Bees Nest Ridge sites 1 and 2, Yellow Rock Rd sites 2 and 3) were composed of all relatively young individuals, with all plants less than 2 m, in contrast to the maximum height for the species of 6–7 m.

### Morphological analysis of species boundaries

Morphological clustering via UPGMA and ordination by semi-strong hybrid multidimensional scaling separated the 48 OTUs scored for morphological characters into three distinct groups: A) *P. cineolifera*; B) a mixed group of *P.* sp. Hawkesbury *P*. sp. Olney State Forest and *P.* sp. Oxley Wild Rivers National Park and C) a mixed group of *P. lanceolata* and *P. ovalifolia*. The top five Kruskal–Wallis characters useful for distinguishing these groups included calyx and pedicel characters (Table 2). Morphometric ordination analysis produced a low stress value (0.0839) indicating an excellent representation of the data in reduced dimensions. The characters contributing most to the discrimination of these groups (Table 2) were four of the five most important in the UPGMA.

**Table 2.** Characters contributing strongly to the distinction of three groups in the morphometric ordination and in the morphometric classification (Fig. 2). For the morphometric ordination, *r*^2^ values ≥ 0.8 are bolded. For the morphometric classification the top five Kruskal–Wallis (KW) values are bolded. A = *Prostanthera cineolifera*; B = *P.* sp. Hawkesbury + *P*. sp. Olney State Forest + *P.* sp. Oxley Wild Rivers National Park; C = *P. lanceolata* + *P. ovalifolia* + *P*. sp. Mt Marsh. The a_1_ axis of the pedicel describes the portion between the bracteoles and the base of the pedicel

| Character (units) | $r^2$ | KW | A | B | C |
| --- | --- | --- | --- | --- | --- |
| Calyx lobes, shape in bud | **0.902** | **37.4** | lobes appressed | indeterminate | lobes reflexed |
| Prophylls, persistent or caducous | **0.859** | 25.1 | persistent | caducous | persistent |
| Hairs on calyx rim, density | **0.849** | **35.7** | usually none - sparse | usually dense | ranges from none to dense |
| Leaf length (mm) | **0.843** | 19.4 | 16–30 | 18.5–36 | 10.5–28 |
| $a_1$ axis length<sup>a</sup> (mm) | 0.792 | **34.2** | 1.6–3.1 | 0–1.7 | 0.5–2.0 |
| $a_1$ axis + anthopodium length<sup>a</sup> (mm) | 0.784 | **33.7** | 1.9–4.4 | 0.5–1.9 | 1.4–2.8 |
| Hairs on the calyx rim length (mm) | 0.658 | **34.1** | 0–0.06 | 0.04–0.14 | 0–0.06 |

<sup>a</sup> pedicel characters

The putative population of *P. cineolifera* at Tabbimoble Creek (TB1) in northeastern New South Wales clustered with the *P. lanceolata* and *P. ovalifolia* group in both the morphometric phenogram (C in Fig. 2a) and the morphometric ordination (C in Fig. 2b). The immature Pillar Valley samples were not included. In Group B, the three samples from the one population in Olney State Forest form a separate branch; the second branch is mixed (Fig. 2a). In Group C, each of the three main branches are mixed samples of *P. lanceolata* and *P. ovalifolia* (Fig. 2a).

**Fig. 2.**
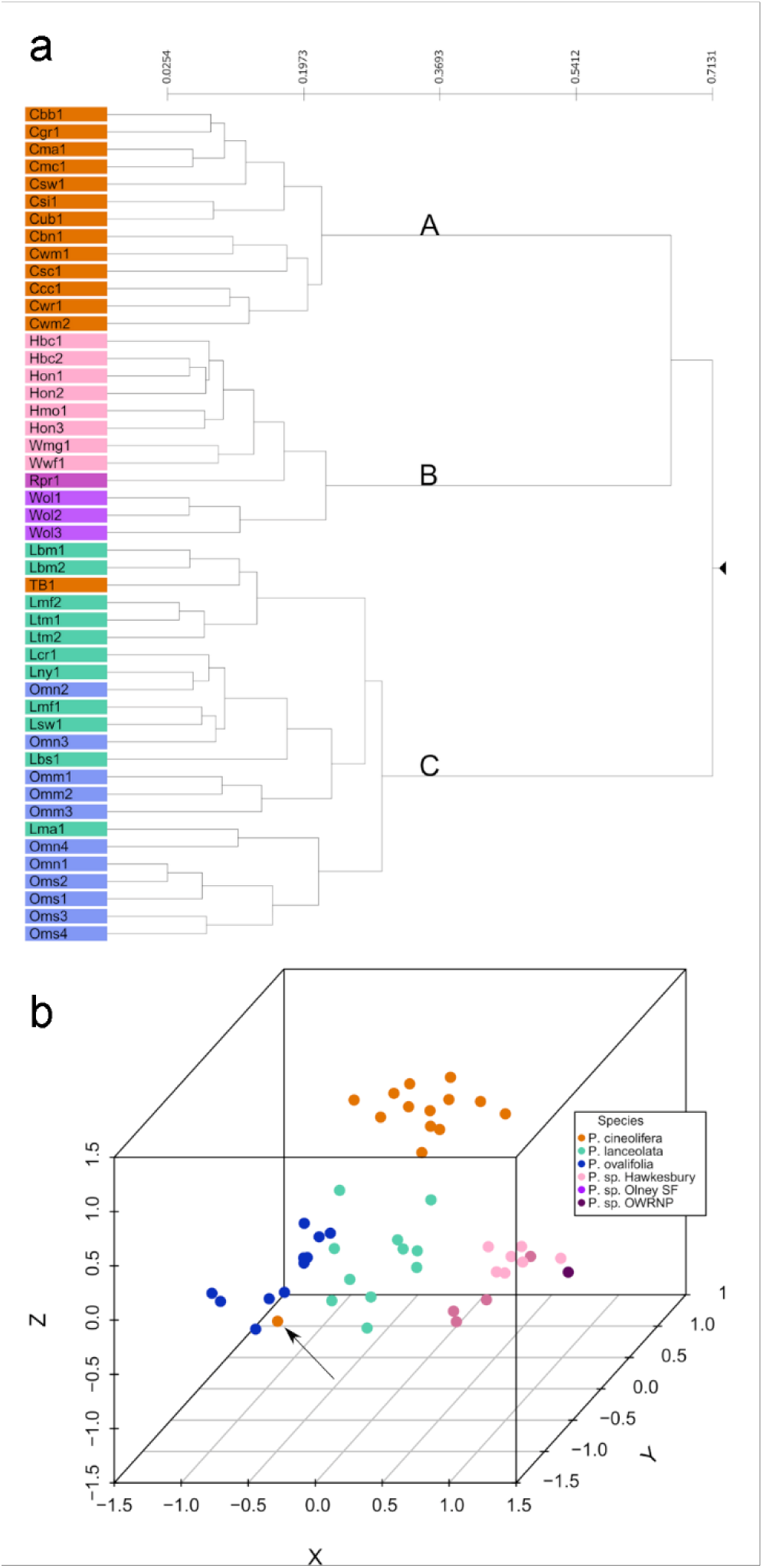
(a) Phenogram from morphometric classification of 48 operational taxonomic units (OTUs) showing three groups. A *Prostanthera cineolifera;* B a mixed group of *P.* sp. Hawkesbury, *P*. sp. Olney State Forest and *P.* sp. Oxley Wild Rivers National Park; and C a mixed group of *P. lanceolata* and *P. ovalifolia* including the specimen from Tabbimoble Creek (TB1) previously determined as *P. cineolifera*. See Table S1 for details of vouchers and OTU codes. (b) Morphometric ordination plot of semi-strong hybrid multidimensional scaling results. The Tabbimoble Creek specimen (TB1) is arrowed

### Molecular analysis of species boundaries

The neighbour-joining tree based on molecular data for all Clade J samples showed distinct clusters of samples (Fig. 3) in concordance with the three clusters of the morphometric classification and morphometric ordination (Fig. 2). All samples of *P. cineolifera* clustered together in Clade A. The populations assigned *a priori* to *P*. sp. Hawkesbury, *P*. sp. Olney State Forest and *P.* sp. Oxley Wild Rivers National Park—represented here as ‘Garabaldi Homestead’—form Clade B. Consequently, ‘the *P.* sp. Hawkesbury group’ refers hereinafter to *P*. sp. Hawkesbury, *P*. sp. Olney State Forest and *P.* sp. Oxley Wild Rivers National Park. Clade C includes all individuals assigned *a priori* to *P. lanceolata* or *P. ovalifolia* plus *P*. sp. Mt Marsh and the ‘unknown’ specimens from Tabbimoble Creek and Pillar Valley that had been previously determined as *P. cineolifera*. Within Clade C, Clade D and Clade E correspond to geographic distribution but not phenology. All Clade D samples were collected north of the border between Queensland and New South Wales—the McPherson Range— while all Clade E samples were collected south of the McPherson range. When used hereinafter, the *P. ovalifolia/lanceolata* group refers to *P. ovalifolia, P. lanceolata* and *P*. sp. Mt Marsh.

**Fig. 3.**
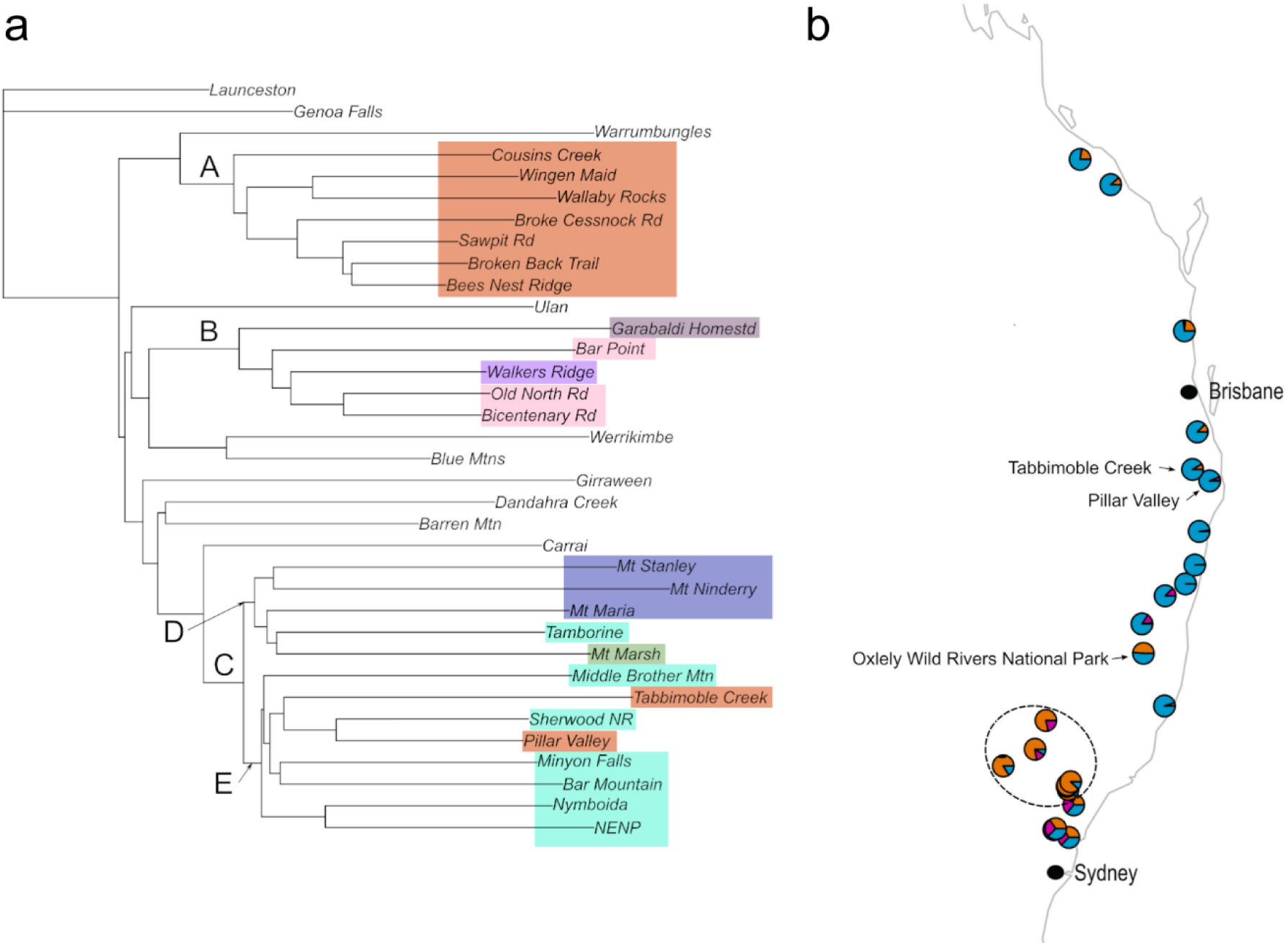
(a) Neighbour-joining tree of molecular data for samples from the corrected set of Clade J taxa. Branch names are sampling locations. Distances are based on 12,107 loci across 144 individuals. A = *Prostanthera cineolifera*, B = *P*. sp. Hawkesbury, *P*. sp. Olney State Forest and *P.* sp. Oxley Wild Rivers National Park and C = *P. lanceolata*, *P. ovalifolia*, *P.* sp. Mt Marsh and the specimens from Tabbimoble Creek and Pillar Valley that had been previously determined as *P. cineolifera*. (b) Non-hierarchical clustering with spatial model in ConStruct (Bradburd *et al*. 2018) of *P. cineolifera, P. lanceolata, P. ovalifolia, P.* sp. Hawkesbury, *P*. sp. Olney State Forest, *P.* sp. Mt Marsh, *P.* sp. Oxley Wild Rivers National Park and the populations from Tabbimoble Creek and Pillar Valley previously determined as *P. cineolifera*. Pies represent estimated proportions of ancestry derived from three layers (see also Figures S7, S8, S9). Dashed ellipse surrounds sampling locations for *P. cineolifera*

Principal coordinate analysis of molecular data for the focal populations (Fig. 4a) identified the same three major clusters as the morphological data. Importantly, all samples that had been determined to be *P. cineolifera* based on morphology clustered together, as did the *P.* sp. Hawkesbury group and the *P. ovalifolia/lanceolata* group, which included the populations in northeastern New South Wales that had been previously identified as *P. cineolifera* and here regarded as unknown. However, further resolution of genetic structure within groups was also possible. PCA of molecular data for the *P. cineolifera* group identified four clusters corresponding to the geographic location of collection site, with all collections in the Pokolbin area forming a tight cluster (Fig. 4b). In the *P.* sp. Hawkesbury group, PCA identified four clusters, mainly corresponding to the geographic location (Fig. 4c). Principal coordinate analysis of molecular data for the *P. ovalifolia/lanceolata* group identified four clusters. Populations flowering in November (Mt Stanley, Mt Maria and Mt Ninderry) formed three separate clusters while populations flowering in September formed one cluster (Fig. 4d).

**Fig. 4.**
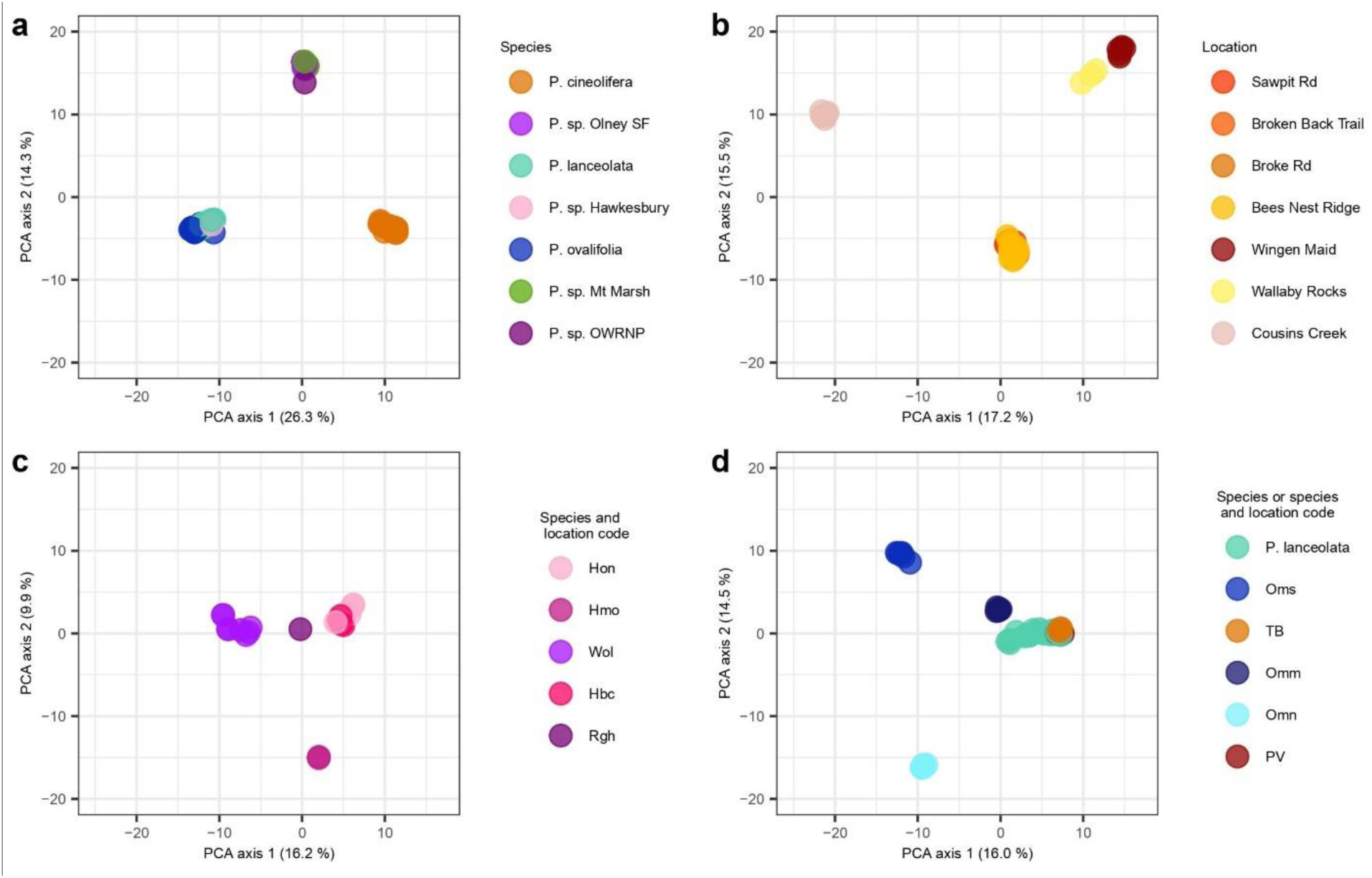
Patterns of genetic structure identified by principal coordinate analysis (PCA) of SNP data. (a) The corrected Clade J taxa (122 individuals). (b) The *Prostanthera cineolifera* taxa of 53 individuals. (c) A mixed group of *P.* sp. Hawkesbury (Hbc, Hmo and Hon), *P*. sp. Olney State Forest (Wol) and *P.* sp. Oxley Wild Rivers National Park (Rgh). (d) A mixed group of *P. lanceolata* and *P. ovalifolia* (Omm, Omn and Oms) including the samples from the populations at Tabbimoble Creek (TB) and Pillar Valley (PV) that had been previously determined as *P. cineolifera.* Percentage of genetic variation explained by each of the PCA axes is provided in parentheses

In ConStruct, the spatial models were best supported up to K=5, and the most stable results with high predictive accuracy were obtained at K=3. Strong differentiation was observed between *P. cineolifera* and the group consisting of *P. lanceolata* and *ovalifolia* (Fig. 3b), but no differentiation was observed between the latter for values of K up to 5 (Fig. S8). The genetic composition of the *P.* sp. Hawkesbury group differed substantially from that of *P. cineolifera* and *P. lanceolata*/*ovalifolia*

The pairwise *F_ST_* values between *P. cineolifera* and *P. lanceolata*, *P. ovalifolia*, *P.* sp. Hawkesbury and *P*. sp. Olney State Forest were all greater than 0.4, indicating pronounced genetic differentiation of *P. cineolifera* from the other species (Table 3). Concordant with the molecular neighbour joining tree (Fig. 3a), there was moderate genetic differentiation between *P. lanceolata* and *P. ovalifolia* (*F_ST_* = 0.19) and between *P*. sp. Hawkesbury and *P*. sp. Olney State Forest (*F_ST_* = 0.14) (3).

**Table 3.** Pairwise *F_ST_* among *Prostanthera* study species. For all pairs, *p* < 0.001 based on 1000 random permutations

|  | *Prostanthera*<br>*cineolifera* | *P. lanceolata* | *P. ovalifolia* | *P. sp. Hawkesbury* |
| --- | --- | --- | --- | --- |
| *P. lanceolata* | 0.439 |  |  |  |
| *P. ovalifolia* | 0.484 | 0.190 |  |  |
| *P. sp. Hawkesbury* | 0.442 | 0.416 | 0.463 |  |
| *P. sp. Olney State*<br>*Forest* | 0.462 | 0.422 | 0.474 | 0.140 |

### Population structure within P. cineolifera

Within *P. cineolifera*, molecular PCA indicated clusters according to the geographic location of collection site, with all collections in the Broken Back Range/Pokolbin area forming a tight cluster (Fig. 4b and extra PCA Analyses in Supplementary Information). Inbreeding was low or absent in most populations, with the exception of Bees Nest Ridge 3, which had an inbreeding coefficient of 0.135 (Table 4). The population sampled at Bees Nest Ridge 3 had the highest gene diversity (*H_E_* = 0.249;), while the more isolated Wingen Maid and Cousins Creek populations had the lowest gene diversity (*H_E_* = 0.15 and *H_E_* = 0.199, respectively).

**Table 4.** Genetic summary statistics for populations of *Prostanthera cineolifera. N* = number of individuals, *H_O_* = observed heterozygosity, *H_E_* = expected heterozygosity, *F* = inbreeding coefficient

| Population | $N$ | $H_o$ | $H_E$ | $F$ |
| --- | --- | --- | --- | --- |
| Bees Nest Ridge 1 | 6 | 0.228 | 0.218 | -0.043 |
| Bees Nest Ridge 3 | 6 | 0.206 | 0.249 | 0.135 |
| Bees Nest Ridge 4 | 7 | 0.23 | 0.243 | 0.038 |
| Broken Back Trail | 6 | 0.223 | 0.229 | 0.016 |
| Cousins Creek | 8 | 0.182 | 0.199 | 0.073 |
| Sawpit Road | 6 | 0.235 | 0.245 | 0.027 |
| Wingen Maid | 6 | 0.149 | 0.15 | -0.008 |

The global *F_ST_* was 0.274, indicating high genetic differentiation among populations within *P. cineolifera*. Pairwise *F_ST_* ranged from 0.025 (little genetic differentiation) between the Bees Nest Ridge populations 3 and 4 to 0.537 (pronounced genetic differentiation) between the Cousins Creek and Wingen Maid populations (Table 5).

**Table 5.** Pairwise F_ST_ among the populations of *P. cineolifera*, where N > 5. Pairwise F_ST_ among the populations of *P. cineolifera*, where N > 5. For all pairs, p < 0.001 based on 100 random permutations. For ease of interpretation, cells are shaded according to Freeland et al. (2011): values less than 0.05 are considered low (pale blue), those between 0.05 and 0.25 are considered moderate (mid blue), and those greater than 0.25 are considered pronounced (darker blue)

|  | Bees Nest Ridge 1 | Bees Nest Ridge 3 | Bees Nest Ridge 4 | Broken Back Trail | Sawpit Rd | Cousins Creek |
| --- | --- | --- | --- | --- | --- | --- |
| Bees Nest Ridge 3 | 0.086 |  |  |  |  |  |
| Bees Nest Ridge 4 | 0.092 | 0.025 |  |  |  |  |
| Broken Back Trail | 0.124 | 0.056 | 0.067 |  |  |  |
| Sawpit Road | 0.104 | 0.038 | 0.046 | 0.077 |  |  |
| Cousins Creek | 0.396 | 0.354 | 0.352 | 0.376 | 0.349 |  |
| Wingen Maid | 0.433 | 0.378 | 0.376 | 0.409 | 0.379 | 0.537 |

The geographically close Broken Back Range populations of Bees Nest Ridge 1, Bees Nest Ridge 3, Bees Nest Ridge 4, Broken Back Trail and Sawpit Rd occur within an area of c. 8 km^2^. For these populations, *F_ST_* was less than 0.125 between all pairs. Greater divergence (*F_ST_* > 0.34) occurred between the Broken Back Range populations and the geographically distant populations of Cousins Creek and Wingen Maid.

Many of the pairwise *F_ST_* values between populations of *P. cineolifera* (Table 5) were similar to the highest of the between-species values (Table 3), for example between *P. cineolifera* and *P. lanceolata*, *F_ST_* = 0.439, and much higher than the between-species divergence of *P. lanceolata* and *P. ovalifolia* (*F_ST_* = 0.19) and for *P.* sp. Hawkesbury and *P.* sp. Olney State Forest (*F_ST_* = 0.14).

Model-based clustering with sNMF also detected strong population structure within *P. cineolifera.* Cross-entropy values had a minimum at *K* = 3 supporting a model with three genetic clusters/ancestral populations (Fig. 5a): Wallaby Rocks and Wingen Maid combined as one cluster, with the Pokolbin area and Cousins Creek as two separate clusters (Fig. 5c), as in the plot of molecular PCA Axis 2 vs Axis 1 (Fig. 4b).

**Fig. 5.**
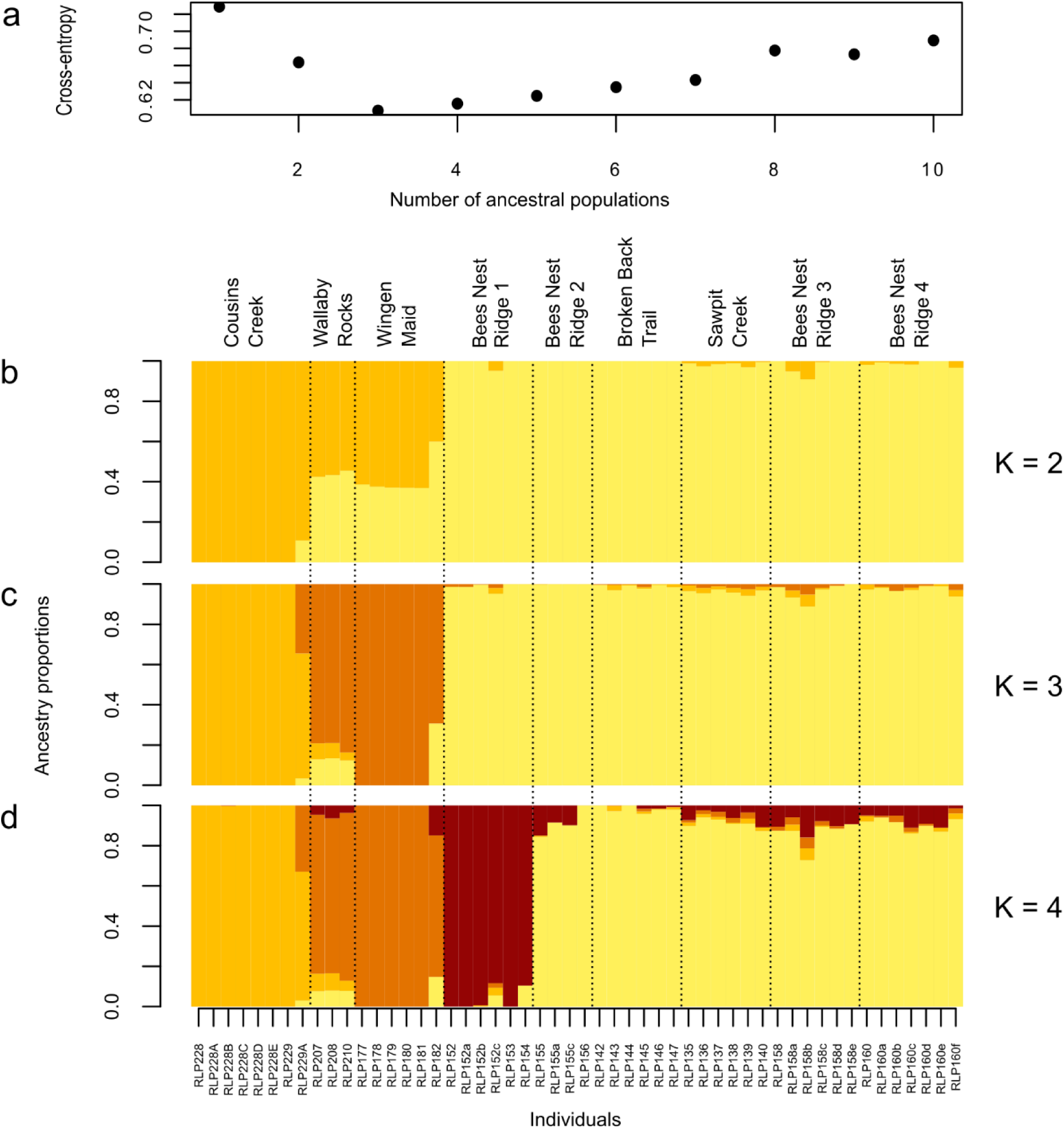
Individual ancestry proportions from model-based clustering with sNMF of the *Prostanthera cineolifera* individuals, assuming there are 2–4 genetic clusters/ancestral populations. (a) Cross entropy values. These values have a minimum at *K* = 3 indicating three genetic cluster/ancestral populations are represented in the data. For K = 2 and K = 3, all 20 replicates indicated clusters as in (b) and (c) respectively. For K = 4, 26 of 30 replicates indicated clusters as shown in (d)

Further model-based clustering provided informative distinctions between geographic sampling locations, indicating the presence of population substructure at relatively fine scales (Fig. 5d, S21, S22). For 26 of 30 replicates for *K* = 4, the southeastern most population on Bees Nest Ridge, Bees Nest Ridge 1, forms a fourth cluster, distinct from the cluster comprised of Sawpit Rd, Broken Back Trail and the other three populations on Bees Nest Ridge (Fig. 5d. Concordant with their very low pairwise *Fst* values (Table 5), Sawpit Rd, Bees Nest Ridge 3 and Bees Nest Ridge 4 were consistently clustered together for *K* < 8 (Figs S21, S22). The three lowest pairwise *Fst* values were observed between three populations (between Bees Nest Ridge 3 and Bees Nest Ridge 4: *Fst* = 0.023, between Bees Nest Ridge 3 and Sawpit Rd: *Fst* = 0.038 and between Bees Nest Ridge 4 and Sawpit Rd: *Fst* = 0.046).

## Discussion

Our study aimed to and to establish the species boundaries of *P. cineolifera* with respect to populations of *Prostanthera* sometimes determined to be *P. cineolifera* or *P. ovalifoilia.* Analyses of both the morphological and molecular data returned the same three distinct groups: A) *P. cineolifera*; B) the *P*. sp. Hawkesbury group, a mixed group of *P.* sp. Hawkesbury, *P*. sp. Olney State Forest and *P.* sp. Oxley Wild Rivers National Park; and C) the *P. lanceolata*/*ovalifolia* group, a mixed group of *P. lanceolata*, *P. ovalifolia* and *P.* sp. Mt Marsh and ‘unknown’ populations of interest in northeastern New South Wales.

### Geographic extent of *P. cineolifera*

Analyses of both the morphological and molecular data supported the conclusion that *P. cineolifera* is more geographically restricted than previously thought, but nevertheless includes populations previously determined to be either *P. discolor* or *P. ovalifolia.* The taxonomic concept of *P. cineolifera* as a morphologically, genetically, geographically, chemically (Sadgrove *et al*. 2020) and ecologically distinct species is supported. We used multiple methods to assess species boundaries in the study group. Patchy sampling may result in apparently discrete population structure, purely due to the presence of isolation-by-distance (IBD) among continuous or connected subpopulations, rather than species boundaries (Pritchard *et al*. 2000; Tapper *et al*. 2014). The IBD pattern in *P. cineolifera*, and more broadly within the study group, suggested that it should be accounted for when testing for discrete population structure, and *conStruct* models incorporating spatial autocorrelation consistently performed better in cross-validation than did nonspatial models (Fig. S7). Clear distinctions exist between *P. cineolifera*, and the other two major groups, but the differentiation between *P. lanceolata* and *P. ovalifolia* could be explained by the underlying spatial autocorrelation due to IBD. This result was consistent with morphometric analysis, which did not separate *P. lanceolata* and *P. ovalifolia*.

The northeastern New South Wales specimens from Tabbimoble Creek and Pillar Valley (Fig. 1b), which have sometimes been determined as *P. cineolifera*, correspond to descriptions of the *P. lanceolata* and *P. ovalifolia* group, and cluster with other specimens determined by us to be *P. lanceolata* or *P. ovalifolia* in the morphometric and molecular analyses. Our results indicate that these populations belong to the *P. lanceolata*/*ovalifolia* group and are not *P. cineolifera*. This confusion may be avoided in the future by updating a couplet in the PlantNet key (Conn 1993) to use calyx characters rather than the currently used indumentum characters.

The populations in the Hawkesbury region that have sometimes been determined to be *P. cineolifera* were represented in the analysis by Wmg1 (NE 49860). In analyses of both the morphological and molecular data, Wmg1 clustered with the Lower Hawkesbury samples that are morphologically similar to *P*. sp. Hawkesbury with its persistent hairy prophylls approximately 2–3 mm long, recurved calyx lobes and a compact inflorescence. This cluster, which included P. sp. Olney State Forest and *P.* sp. Oxley Wild Rivers National Park, is considered a separate entity and will be the subject of a separate paper.

*Prostanthera cineolifera*, distinguished by stem indumentum, calyx features, bud shape and a loose, open inflorescence is distributed in the Sydney Basin and Nandewar Bioregions (IBRA v7.0) (Department of Climate Change Energy the Environment and Water 2020). The two populations from Manilla and Upper Moore Ck (Tamworth) in the Nandewar Bioregion appear to belong to *P. cineolifera* on the basis of morphology, but they have not been recorded since historical herbarium collections were made in 1913 and 1933 respectively.

One area of uncertainty remains, however. Populations of *Prostanthera* in the West Wyalong to Wagga Wagga districts of New South Wales that were originally described by Bentham (1834) as *P. atriplicifolia* A.Cunn. ex *Benth* appear somewhat morphologically similar to *P. cineolifera.* More work is underway to ascertain if they are a separate entity or *P. cineolifera*.

### Genetic relationships and population structure of *P. cineolifera*

Population structure within *P. cineolifera* was expected, as our sampling localities for *P. cineolifera* are up to 100 km apart. We detected substantial structure at multiple scales, using diverse analytical techniques. Ordination and non-hierarchical cluster analysis identified three main clusters, corresponding to geographic areas. At that scale, *F_ST_* was similar to or greater than that between species in this group. For example, *F_ST_* = 0.537 between Cousins Creek and Wingen Maid, compared with *F_ST_* = 0.439 between *P. cineolifera* and *P. lanceolata*. In a perennial shrub, this likely reflects a long period of genetic isolation, suggesting that the patchy distribution of the species results from a combination of historical biogeographical processes and specialized soil requirements, rather than fragmentation due to agricultural intensification.

Differentiation among individual sampling sites was also seen at finer scales, including between neighbouring populations on Bees Nest Ridge that were only c. 300 m apart. The strong genetic structure within *P. cineolifera* indicates a lack of migration between populations and the most isolated known site (Wingen Maid and by association Wallaby Rocks) has the lowest genetic diversity (*H_E_* = 0.15). The genetic structure is partly associated with geographic separation. *P. cineolifera* is probably pollinated by beetles and flies (Wilson *et al*. 2017), and although some beetle pollinators fly up to 70 m between plants (Englund 1993), insect pollination and gravity dispersed seeds do not favour long distance gene flow. This limited gene flow would contribute to the very high *F_ST_* estimates between samples such as Cousins Creek and Wingen Maid (*F_ST_* = 0.537), and to the moderate *F_ST_* estimate between Broken Back Trail and Bees Nest Ridge 1 (*F_ST_* = 0.124). Model-based clustering with sNMF also detected genetic structure at a range of scales (Figs 5, S21, S22).

While there is good evidence to support at least three distinct management units within *P. cineolifera*, it is not known exactly how many populations remain unsampled in remote patches of suitable habitat between localities. We now know of several more recently-discovered populations in the Pokolbin area that were not sampled for molecular analysis: within the Singleton Military Area (SMA), five populations on Yellow Rock Road, another population on Broken Back Trail and several populations in the Cedar Creek biodiversity offset, now Cedar Creek Nature Reserve (Bell 2017). A large population was recently discovered in Goulburn River National Park, Nevertheless, it is clear from our fieldwork that the distribution of *P. cineolifera* is highly patchy, rather than continuous. While our population sampling was exhaustive based on our knowledge at the time, judging from satellite imagery (for example https://maps.six.nsw.gov.au/), there are other areas that likely contain other populations of *P. cineolifera*.

### Conservation status of *P. cineolifera*

Restricted gene flow can have negative consequences for populations, including inbreeding and decreases in genetic diversity and hence a decrease in adaptive potential of the population (Ellstrand and Elam 1993; Reed and Frankham 2003; Garant *et al*. 2007; Markert *et al*. 2010). However, inbreeding coefficients did not seem to be related to population size, but rather demography. Bees Nest Ridge 3 (population approximately 320) had a relatively high inbreeding coefficient of (*F* = 0.135) unlike Bees Nest Ridge 1 (population approximately 30) which had an inbreeding coefficient of -0.043. These populations differ in age structure, with Bees Nest Ridge 1 being a relatively young population lacking large old shrubs, whereas Bees Nest Ridge 3 had a more even distribution of growth stages. As all plants in the young population are adjacent to the road, we suspect that recruitment of the relatively young populations has been triggered by road building or maintenance, either through introduction of seeds by the grader or by stimulation of the soil seed bank by disturbance. Although no obvious signs of inbreeding depression were observed at Bees Nest Ridge 3, further monitoring may be warranted.

Genetic diversity is needed for populations to adapt to changes in the environment, and gene flow is an important source of genetic variation, particularly when other populations possess potentially adaptive variants (Sexton *et al*. 2011). The predicted global warming of 2–7°C by 2100 (Stott and Kettleborough 2002; O’Neill *et al*. 2016) will place smaller isolated populations with lower genetic diversity, such as the populations of *P. cineolifera* at Wingen Maid and Wallaby Rocks, at greater risk of extinction (Aitken *et al*. 2008). While these isolated populations may have evolved low rates of inbreeding depression or maintain sufficiently large populations to avoid it, adaptive capacity may be limited without historical or recent connectivity. The Wingen Maid population is of concern, as it has low genetic diversity (as measured by expected heterozygosity) in comparison with other populations of the species, despite a relatively healthy population size and structure. Further study of the Wingen Maid population, as well as several small populations, may be needed to determine whether the fitness of the population is vulnerable to changes in precipitation and temperature regimes. The very small population (c. 10) at Wallaby Rocks is of particular concern. This population is genetically similar to the Wingen Maid population but much smaller.

The total known census size of *P. cineolifera* is at least 5,200 plants (Table 1), with the majority in the Pokolbin area. Except for the small populations on Broke Road (3) and at Wallaby Rocks, all populations are multi-aged, although several sites on Bees Best Ridge and Yellow Rock Rd lack large old shrubs and are likely to be the result of road building or maintenance. *Prostanthera cineolifera* is currently listed as Vulnerable under both Commonwealth and New South Wales legislation. While the demographic information, genetic data and estimated population sizes of *P. cineolifera* suggest that the species is not in immediate danger of extinction, the apparent lack of genetic connectivity and signs of reduced genetic diversity or inbreeding do raise concern about conservation of certain populations. Although the species is morphologically consistent across the range that we have determined here, it seems likely that unique genetic diversity would be lost if any populations become extinct. Indeed, the apparent disappearance of populations in the Nandewar Bioregion may have already compromised the evolutionary potential of the species.

Further work is needed to test and clarify species boundaries in the *P. lanceolata*/*ovalifolia* group with more extensive sampling, particularly for *P. ovalifolia*. There is some evidence of genetic clustering in ordinations based on molecular data, and populations of this group differ in phenology: specimens from north of the Brisbane River flower in November, while the populations from south of the Brisbane River flower in September. However, the morphological boundary is not clear, and ConStruct analysis did not support the boundary when accounting for spatial autocorrelation. This suggests that the morphological and phenological variation may reflect isolation-by-distance in a more-or-less continuously distributed species, rather than two reproductively isolated species. Indeed, Althofer (1978, p. 47) observed that “as the [*P. ovalifolia*] moves southward”, down the New South Wales, “an almost imperceptible change takes place. The leaves become progressively larger, and small indentations become perceptible….”. Additional sampling is needed to characterise the morphological and genetic variation in the group, which may reflect adaptation to climatic and elevational gradients.

## Conclusions

By clarifying the species boundaries of *Prostanthera cineolifera*, this study has shown that putative populations outside of the Hunter Valley belong to other species and are not of conservation concern as unusual isolates of *P. cineolifera*. The northern populations are part of the *P. lanceolata*/*ovalifolia* group, while the populations in the Lower Hawkesbury are distinct from *P. cineolifera* and will be the subject of a separate taxonomic paper. The study has also documented deep genetic divergence among populations of *P. cineolifera*, probably due to long-term isolation, rather than recent fragmentation. Finer-scale subdivision was also observed, likely as a result of isolation-by-distance, consistent with insect pollination and passive seed dispersal. While the *P. cineolifera* population appears robust, certain populations exhibited lower genetic diversity or elevated inbreeding coefficients, although neither was related to population size. This work highlights the genetic consequences of restricted dispersal and isolated populations and identifies a need for further research on the adaptive capacity of such populations.

## Acknowledgements

We thank Saving Our Species (New South Wales Department of Planning and Environment) and University of New England (UNE) for funding; National Parks and Wildlife Service (New South Wales) and Forestry Corporation New South Wales, various Rangers of National Parks; Luke Foster, Robert Gibson, (New South Wales Department of Planning and Environment); Australian Army, Officer in Charge Singleton Military Area; Peter Bussell, Kaye Durham, Elizabeth Gill, Joan Griffiths, Aaron Hedger, David Martin, Hilary Pearl, Judith Roland, Brigitte Stievermann and Bob Ross for assistance with fieldwork; John Nevin and Phil Rose for assistance with propagation of cuttings and repotting of specimens; Penelope Sinclair for assistance with mounting specimens; Margaret Donald for assistance with field work, giving RLP a good grounding in R and for the use of her Leica M60 microscope; Jody McNally of Commonwealth Scientific and Industrial Research Organization (CSIRO) Chiswick for assistance with freeze drying material for DNA extraction; directors and staff of Herbaria AD, BRI, CANB, and NSW, for help during visits; UNE for access to facilities, in particular, the N.C.W. Beadle Herbarium (NE) where this study was based; Ian Simpson, Lindsey Frost and Rhyan Gorman for assisting watering in the glasshouse; Trish Waters, Margaret and Chris Cooper, Jeremy Bruhl and family, and Laurel Arthur and family provided accommodation for RLP on visits to Armidale and Canberra; and Trevor Wilson for assistance at NSW and valuable discussions with RLP.

## Author contributions

Jeremy Bruhl, Rose Andrew and Ian Telford contributed to the study concept and design. Sample collection was performed by Ruth Palsson, Jeremy Bruhl, Rose Andrew and Ian Telford. Analyses were performed by Ruth Palsson with advice from Rose Andrew, Jeremy Bruhl and Ian Telford. The first draft of the manuscript was written by Ruth Palsson and all authors commented on versions of the manuscript. All authors read and approved the final manuscript.

## Data Availability

Morphological data are available in the accompanying Supplementary Information. SNP matrices will be publicly available from the University of New England data repository, DOI XX.XXXXX.

