## Supplementary Materials for "Population genetic structure and range limits of *Prostanthera cineolifera* (Lamiaceae), a vulnerable shrub with a patchy distribution"

Contents

Table S1. Collections used for used for phenetic analysis (P) and/or molecular analyses (G) ...2
BRI AQ0336457 ...6
Table S2. Character list used for the morphological study ...8
Table S3. Morphological data used in PATN analysis ...10
Extra morphometric ordination plots – includes Figs S3 and S4 ...12
Table S4. Details of samples sent to DArT ...15
Technical duplicates including Table S5 ...22
Anomalous specimens ...24
Table S6. Filters used and the resulting numbers of individuals and loci ...25
Table S7. Samples removed during the filtering process due to low call rates ...26
Table S8. Basic summary statistics for the main species of the project ...26
conStruct analysis including Figs S7, S8, S9 ...27
Extra PCA analyses ...30
Figs S21, S22 LEA plots ...36
References ...38

**Table S1 Collections used for used for phenetic analysis (P) and/or molecular analyses (G)**

Bolded OTU code indicates a type specimen. Ck = Creek, NP = National Park, NSW = New South Wales, Qld = Queensland, Rd = Road, SF = State Forest. Herbarium codes follow Index Herbariorum (see http://sweetgum.nybg.org/ih/ accessed 14 December 2018).

| Putative taxon | OTU code | DArtT code | Analyses | Location | Collector(s) & number | Herbarium & accession no. |
|---|---|---|---|---|---|---|
| *P. cineolifera* | Cma1 | | P | Manilla, NSW | *B.S. Morse s.n.* | NSW 134412 |
| *P. cineolifera* | Cmc1 | | P | Upper Moore Creek, Tamworth, NSW | *E. Wyndham s.n.* | NSW 134411 |
| *P. cineolifera* | | RLP181-1 | G | Wingen Maid, NSW | *R.L. Palsson 181 et al.* | n/a |
| *P. cineolifera* | | RLP182-1 | G | Wingen Maid, NSW | *R.L. Palsson 182 et al.* | n/a |
| *P. cineolifera* | | RLP177-1 | G | Wingen Maid, NSW | *R.L. Palsson 177 et al.* | NE 106344 |
| *P. cineolifera* | | RLP180-1 | G | Wingen Maid, NSW | *R.L. Palsson 180 et al.* | NE 106347 |
| *P. cineolifera* | Cwm2 | RLP179-1 | P, G | Wingen Maid, NSW | *R.L. Palsson 179 et al.* | NE 106346 |
| *P. cineolifera* | Cwm1 | RLP178-1 | P, G | Wingen Maid, NSW | *R.L. Palsson 178 et al.* | NE 106345 |
| *P. cineolifera* | Csc1 | | P | Scone, NSW | *R.H. Cambage s.n.* | NSW 227969 |
| *P. cineolifera* | Cwr1 | | P | Wallaby Rocks, NSW | *N.J. Sadgrove 291* | NE 105284 |
| *P. cineolifera* | | RLP207-1 | G | Wallaby Rocks, NSW | *R.L. Palsson 207* | NE 106072 |
| *P. cineolifera* | | RLP208-1 | G | Wallaby Rocks, NSW | *R.L. Palsson 208* | NE 106073 |
| *P. cineolifera* | | RLP210-1 | G | Wallaby Rocks, NSW | *R.L. Palsson 210* | NE 106074 |
| *P. cineolifera* | Cgr1 | | P | Goulburn River NP, NSW | *G. Ryan s.n.* | NSW 227965 |
| *P. cineolifera* | Ccc1 | | P | Cousins Creek, Wollemi NP, NSW | *S.A.J. Bell s.n.* | NSW 424206 |
| *P. cineolifera* | | RLP228-1 | G | Cousins Creek, Wollemi NP, NSW | *R.L. Palsson 228 & L.M.C. Foster* | NE 108826 |
| *P. cineolifera* | | RLP229-1 | G | Cousins Creek, Wollemi NP, NSW | *R.L. Palsson 229 & L.M.C. Foster* | NE 108827 |
| *P. cineolifera* | | RLP148-2 | G | Broke Rd, Pokolbin, NSW | *R.L. Palsson 148 et al.* | NE 106407 |
| *P. cineolifera* | Cbb1 | | P | Broken Back Range, Broken Back Trail, NSW | *L.M.C. Foster 1a* | n/a |
| *P. cineolifera* | Cbn1 | RLP155-1 | P, G | Broken Back Range, Bees Nest Rd, NSW | *R.L. Palsson 155 et al.* | NE 106411 |
| *P. cineolifera* | | RLP152-1 | G | Broken Back Range, Bees Nest Rd, BNR_1, NSW | *R.L. Palsson 152 et al.* | NE 106409 |
| *P. cineolifera* | | RLP154-2 | G | Broken Back Range, Bees Nest Rd, BNR_1, NSW | *R.L. Palsson 154 et al.* | NE 106412 |
| *P. cineolifera* | | RLP153-2 | G | Broken Back Range, Bees Nest Rd, BNR_1, NSW | *R.L. Palsson 153 et al.* | NE 106413 |

| **Putative taxon** | **OTU code** | **DArtT code** | **Analyses** | **Location** | **Collector(s) & number** | **Herbarium & accession no.** |
|---|---|---|---|---|---|---|
| *P. cineolifera* | | RLP156-2 | G | Broken Back Range, Bees Nest Rd, BNR_2, NSW | *R.L. Palsson 156 et al.* | NE 106414 |
| *P. cineolifera* | | RLP158-2 | G | Broken Back Range, Bees Nest Rd, BNR_3, NSW | *R.L. Palsson 158 et al.* | NE 106331 |
| *P. cineolifera* | | RLP160-1 | G | Broken Back Range, Bees Nest Rd, BNR_4, NSW | *R.L. Palsson 160 et al.* | NE 106008 |
| *P. cineolifera* | **Csi1-T** | | P | Siberia, Broke, NSW | *C.H. Cheesborough s.n.* | NE 106007 |
| *P. cineolifera* | | RLP142-1 | G | Broken Back Range, Broken Back Trail, NSW | *R.L. Palsson 142 et al.* | NE 106275 |
| *P. cineolifera* | | RLP143-1 | G | Broken Back Range, Broken Back Trail, NSW | *R.L. Palsson 143 et al.* | NE 105883 |
| *P. cineolifera* | | RLP144-1 | G | Broken Back Range, Broken Back Trail, NSW | *R.L. Palsson 144 et al.* | NE 105880 |
| *P. cineolifera* | | RLP146-2 | G | Broken Back Range, Broken Back Trail, NSW | *R.L. Palsson 146 et al.* | NE 105873 |
| *P. cineolifera* | | RLP147-1 | G | Broken Back Range, Broken Back Trail, NSW | *R.L. Palsson 147 et al.* | NE 105874 |
| *P. cineolifera* | | RLP145-1 | G | Broken Back Range, Broken Back Trail, NSW | *R.L. Palsson 145 et al.* | NE 105877 |
| *P. cineolifera* | Cub1 | | P | Upper Bellbird, NSW | *L.M.C. Foster s.n.* | NE 105876 |
| *P. cineolifera* | Csw1 | RLP139-1 | P, G | Broken Back Range, Sawpit Rd, NSW | *R.L. Palsson 139 & M.R. Donald* | NE 106000 |
| *P. cineolifera* | | RLP135-1 | G | Broken Back Range, Sawpit Rd, NSW | *R.L. Palsson 135 & M.R. Donald* | NE 100319 |
| *P. cineolifera* | | RLP136-1 | G | Broken Back Range, Sawpit Rd, NSW | *R.L. Palsson 136 & M.R. Donald* | NE 107489 |
| *P. cineolifera* | | RLP138-1 | G | Broken Back Range, Sawpit Rd, NSW | *R.L. Palsson 138 & M.R. Donald* | NE 106857 |
| *P. cineolifera* | | RLP140-2 | G | Broken Back Range, Sawpit Rd, NSW | *R.L. Palsson 140 & M.R. Donald* | NE 106864 |
| *P. cineolifera* | | RLP137-1 | G | Broken Back Range, Sawpit Rd, NSW | *R.L. Palsson 137 & M.R. Donald* | NE 106213 |
| *P. incisa* | | JJB3532-1 | G | Werrikimbe NP, NSW | *J.J. Bruhl 3532 & F.C. Quinn* | NE 106214 |
| *P. incisa* | Isb1-T | | P | Sydney Basin, NSW | *R. Brown s.n.* | NE 106220 |
| *P. lanceolata* | Ltm3 | | P | Tamborine Mtn, Qld | *H. Curtis s.n.* | BRI AQ0336457 |
| *P. lanceolata* | Ltm2 | RLP107-1 | P, G | Cliff Walk, Tamborine Mtn, Qld | *R.L. Palsson 107 et al.* | NE 106180 |
| *P. lanceolata* | Ltm1 | RLP104-1 | P, G | Cliff Walk, Tamborine Mtn, Qld | *R.L. Palsson 104 et al.* | NE 106182 |
| *P. lanceolata* | Lbm1 | RLP110a-1 | P, G | Bar Mtn, Border Ranges NP, NSW | *R.L. Palsson 110 et al.* | NE 106181 |
| *P. lanceolata* | Lbm2 | | P | Bar Mtn, Border Ranges NP, NSW | *R.L. Palsson 111 et al.* | NE 106207 |
| *P. lanceolata* | Lmf1 | RLP114-1 | P, G | Minyon Falls, Nightcap NP, NSW | *R.L. Palsson 114 et al.* | NE 106206 |
| *P. lanceolata* | Lmf2 | RLP113-1 | P, G | Minyon Falls, Nightcap NP, NSW | *R.L. Palsson 113 et al.* | NE 106203 |
| *P. lanceolata* | Lsw1 | RLP122-1 | P, G | Sherwood NR, NSW | *R.L. Palsson 122 et al.* | NE 103755 |

| **Putative taxon** | **OTU code** | **DArtT code** | **Analyses** | **Location** | **Collector(s) & number** | **Herbarium & accession no.** |
|---|---|---|---|---|---|---|
| *P. lanceolata* | Lny1 | NJS239-1 | P, G | Nymboida River, NSW | *N.J. Sadgrove 239 & I.R. Telford* | NE 105338 |
| *P. lanceolata* | | JJB3563-1 | G | New England NP, NSW | *J.J. Bruhl 3563 et al.* | NE 108059 |
| *P. lanceolata* | Lcr1 | PGW1890-1 | P, G | Carrai Rd (The Castle NR), NSW | *P.G. Wilson 1890 & M.M. Heslewood* | NE 108061 |
| *P. lanceolata* | Lbs1 | | P | Mt Boss SF, NSW | *H. Streimann 8216* | BRI 313225 |
| *P. lanceolata* | | JJB3542-1 | G | Middle Brother Mtn NP, NSW | *J.J. Bruhl 3542* | NE 83318 |
| *P. latifolia* | | IRT13512-2 | G | Tapin Tops, NSW | *I.R. Telford 13512* | NE 104451 |
| *P. latifolia* | | NJS424-2 | G | Tapin Tops, NSW | *N.J. Sadgrove 424 & I.R. Telford* | NE 108063 |
| *P. ovalifolia* | Omw1-T | | P | Mt Westall, Qld | *R. Brown s.n.* | NE 106309 |
| *P. ovalifolia* | Omp1 | | P | Mt Parnassus, Qld | *J. Brushe 2148* | NE 106323 |
| *P. ovalifolia* | Oms1 | RLP195-1 | P, G | Mt Stanley, Qld | *R.L. Palsson 195 et al.* | NE 106314 |
| *P. ovalifolia* | Oms3 | | P | Mt Stanley, Qld | *R.L. Palsson 195a et al.* | NE 106319 |
| *P. ovalifolia* | Oms4 | RLP200-1 | P, G | M Stanley, Qld | *R.L. Palsson 200 et al.* | NE 106322 |
| *P. ovalifolia* | Oms2 | RLP201-1 | P, G | Mt Stanley, Qld | *R.L. Palsson 201 et al.* | NE 106320 |
| *P. ovalifolia* | Omm1 | RLP188-1 | P, G | Mt Maria, Qld | *R.L. Palsson 188 et al.* | NE 107073 |
| *P. ovalifolia* | Omm2 | | P | Mt Maria, Qld | *R.L. Palsson 188d et al.* | NE 104451 |
| *P. ovalifolia* | Omm3 | | P | Mt Maria, Qld | *R.L. Palsson 188a et al.* | NE 108063 |
| *P. ovalifolia* | Omn1 | RLP185-2 | P, G | Mt Ninderry, Qld | *R.L. Palsson 185 et al.* | NE 49860 |
| *P. ovalifolia* | Omn4 | | P | Mt Ninderry, Qld | *P.R. Sharpe 2076* | NE 106385 |
| *P. ovalifolia* | Omn2 | | P | Mt Ninderry, Qld | *R.L. Palsson 184e et al.* | NE 106002 |
| *P. ovalifolia* | Omn3 | RLP184-1 | P, G | Mt Ninderry, Qld | *R.L. Palsson 184 et al.* | NE 106341 |
| *P.* sp. Barren Mtn | | NJS366-2 | G | Barren Mtn, NSW | *N.J. Sadgrove 366 & G.T. Plunkett* | NE 106340 |
| *P.* sp. Barren Mtn | | NJS428-2 | G | Barren Mtn, NSW | *N.J. Sadgrove 428* | NE 106339 |
| *P.* sp. Dandahra Ck | | JJB3515-1 | G | Dandahra Ck, NSW | *J.J. Bruhl 3515 & I.R. Telford* | NE 106212 |
| P. sp. Dandahra Ck | | NJS296-1 | G | Dandahra Ck, NSW | *N.J. Sadgrove 296* | NE 106342 |
| *P.* sp. Hawkesbury | Wmg1 | | P | Mangrove Ck, NSW | *D.S. Gibbons s.n. & I.R. Telford* | NE 49860 |
| *P.* sp. Hawkesbury | Hon3 | | P | Old Northern Rd, Wisemans Ferry, NSW | *M.R. Donald 108 & R.L. Palsson* | NE 106385 |
| *P.* sp. Hawkesbury | Wwf1 | | P | Wisemans Ferry, NSW | *D.W. Shoobridge s.n.* | NE 106002 |

| **Putative taxon** | **OTU code** | **DArtT code** | **Analyses** | **Location** | **Collector(s) & number** | **Herbarium & accession no.** |
|---|---|---|---|---|---|---|
| *P*. sp. Hawkesbury | | RLP176-2 | G | Old Northern Rd, Wisemans Ferry, NSW | *R.L. Palsson 176 et al.* | NE 106341 |
| *P*. sp. Hawkesbury | | MRD105-1 | G | Old Northern Rd, Wisemans Ferry, NSW | *M.R. Donald 105 & R.L. Palsson* | NE 106212 |
| *P*. sp. Hawkesbury | | RLP173-1 | G | Old Northern Rd, Wisemans Ferry, NSW | *R.L. Palsson 173 et al.* | NE 106342 |
| *P*. sp. Hawkesbury | Hon2 | | P | Old Northern Rd, Wisemans Ferry, NSW | *R.L. Palsson 175 et al.* | NE 106340 |
| *P*. sp. Hawkesbury | Hon1 | RLP174-1 | P, G | Old Northern Rd, Wisemans Ferry, NSW | *R.L. Palsson 174 et al.* | NE 106339 |
| *P*. sp. Hawkesbury | Hbc1 | RLP171-2 | P, G | Bicentenary Rd, Webbs Creek, NSW | *R.L. Palsson 171 et al.* | NE 106336 |
| *P*. sp. Hawkesbury | Hbc2 | RLP170-1 | P, G | Bicentenary Rd, Webbs Creek, NSW | *R.L. Palsson 170 et al.* | NE 106337 |
| *P*. sp. Hawkesbury | | RLP172-2 | G | Bicentenary Rd, Webbs Creek, NSW | *R.L. Palsson 172 et al.* | NE 106338 |
| *P*. sp. Hawkesbury | | RLP167-1 | G | Bicentenary Rd, Webbs Creek, NSW | *R.L. Palsson 167 et al.* | NE 106006 |
| *P*. sp. Hawkesbury | | RLP168-1 | G | Bicentenary Rd, Webbs Creek, NSW | *R.L. Palsson 168 et al.* | NE 106355 |
| *P*. sp. Hawkesbury | | RLP169-1 | G | Bicentenary Rd, Webbs Creek, NSW | *R.L. Palsson 169 et al.* | NE 105986 |
| *P*. sp. Hawkesbury | Hmo1 | | P | Bar Point, NSW | *R.L. Palsson 133 & M.R. Donald* | NE 105998 |
| *P*. sp. Hawkesbury | | RLP206-1 | G | Bar Point, NSW | *R.L. Palsson 206* | NE 105999 |
| *P*. sp. Mt Marsh | Lma1 | LMC3770-2 | P, G | Mt Marsh, NSW | *L.M. Copeland 3770 et al.* | NE 106336 |
| *P*. sp. Olney State Forest | Wol3 | RLP164-1 | P, G | Olney SF, NSW | *R.L. Palsson 164 et al.* | NE 106337 |
| *P*. sp. Olney State Forest | Wol1 | RLP166-2 | P, G | Olney SF, NSW | *R.L. Palsson 166 et al.* | NE 106338 |
| *P*. sp. Olney State Forest | | RLP163-1 | G | Olney SF, NSW | *R.L. Palsson 163 et al.* | NE 106006 |
| *P*. sp. Olney State Forest | | RLP165-1 | G | Olney SF, NSW | *R.L. Palsson 165 et al.* | NE 106355 |
| *P*. sp. Olney State Forest | | RLP162-1 | G | Olney SF, NSW | *R.L. Palsson 162 et al.* | NE 105986 |
| *P*. sp. Olney State Forest | Wol2 | RLP161-1 | P, G | Olney SF, NSW | *R.L. Palsson 161 et al.* | NE 105998 |
| *P*. sp. Oxley Wild Rivers National Park | Rpr1 | MFD4027-2 | P, G | Paradise Rocks, NSW | *M.F. Duretto 4027 & T.L. Collins* | NE 105999 |
| Unknown | TB1 | RLP119-1 | P, G | Tabbimoble Ck, NSW | *R.L. Palsson 119 et al.* | NE 105986 |
| Unknown | | RLP120-1 | G | Tabbimoble Ck, NSW | *R.L. Palsson 120 et al.* | NE 105998 |
| Unknown | | RLP121-2 | G | Pillar Valley, NSW | *R.L. Palsson 121 et al.* | NE 105999 |

**BRI AQ0336457**

An herbarium search at the Queensland Herbarium (BRI) revealed a specimen (BRI AQ0336457) determined as *P. ovalifolia* and reportedly collected at Tamborine Mtn in southeast Qld by H. Curtis in 1944. This specimen had the stem indumentum, bud shape and calyx shape typical of *P. cineolifera*. This specimen was included in initial phenetic analyses with the OTU code Ltm3. As expected, in these initial phenetic analyses (Fig S1), this specimen clustered with specimens of *P. cineolifera*.

*Prostanthera* on Tamborine Mtn were expected to be *P. lanceolata*—Tamborine Mtn is the type locality of *P. lanceolata.* All herbarium specimens of *P. lanceolata* or *P. ovalifolia* collected on Tamborine Mtn and on AVH have been examined by RLP or Mike Mathieson at BRI. All specimens except Ltm3 (BRI AQ 336457) correspond to descriptions of *P. lanceolata*. Additional known populations of *Prostanthera* on Tamborine Mtn that are not represented in herbarium collections have been visited by RLP with Judith Roland of Tamborine Mountain Landcare Inc. These populations are consistent with *P. lanceolata*; Ltm3 remains anomalous, but its identity cannot be confirmed without relocating its source population. Currently, we consider this location data recorded on BRI AQ0336457 spurious. It was removed from further analyses.

As far as can be determined, H. Curtis (née Geissmann), a respected naturalist and photographer (Curtis, 2015), submitted about 50 specimens to Australian herbaria (AVH, 2017). Three of these specimens were probably submitted by a different H. Curtis (two marine algae from Tasmania and *Pandanus tectorius* from north Queensland). All but one of other H. Curtis specimens were collected in south-east Queensland. The other was collected from north-east New South Wales. The Curtis diaries may shed light on this issue if they become available for examination.

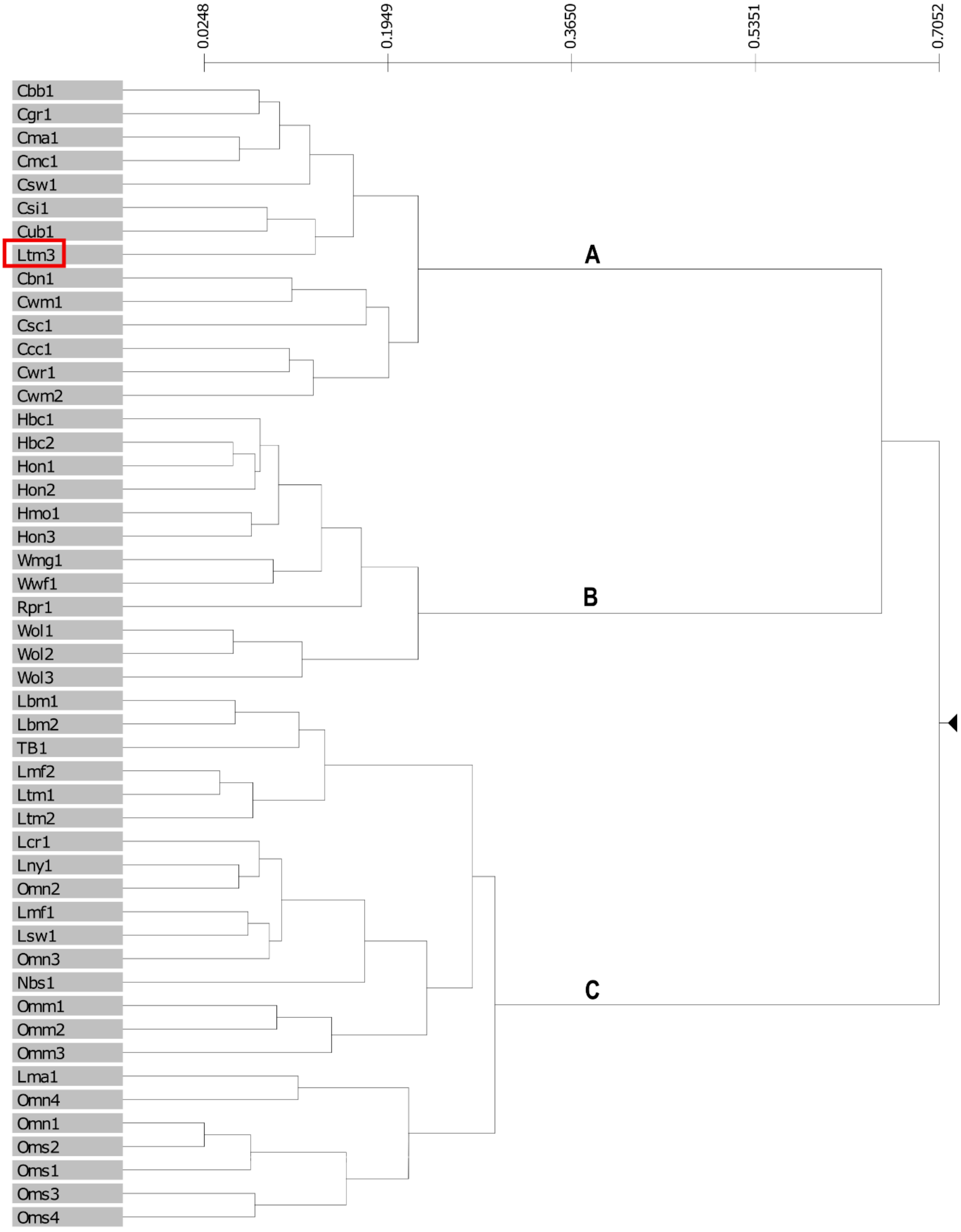


**Fig. S1** Phenogram from morphometric classification of 49 operational taxonomic units (OTUs) showing three groups. A *Prostanthera cineolifera;* B a mixed group of *P.* sp. Hawkesbury, *P.* sp. Olney State Forest and *P.* sp. Oxley Wild Rivers National Park; and C a mixed group of *P. lanceolata* and *P. ovalifolia*. The anomalous H. Curtis specimen (BRI AQ0336457) is boxed. See Table S1 for details of vouchers.

**Table S1 Character list used for the morphological study**

Number in brackets indicates character state.

| | |
|---|---|
| 1 | Hairs on branches (distribution): whether |
| | in rows on the decurrencies (1); |
| | scattered (not in rows) (2); or |
| | distinct rows on the decurrencies and widespread between the decurrencies (3) |
| 2 | Hairs on branches (shape): whether |
| | antrorse (1); or |
| | divergent (2) |
| 3 | Leaf lamina (length): cm long |
| 4 | Leaf lamina (width): cm wide |
| 5 | Leaf widest point (measured from base of lamina): cm wide |
| 6 | Prophylls: whether |
| | persistent (0); or |
| | caducous (1) |
| 7 | Prophylls (length): mm long |
| 8 | [a]Anthopodium plus $a_1$ axis (length): mm long |
| 9 | $a_1$ axis (length): mm long |
| 10 | Second internode from the tip of a flowering branchlet (length): mm long |
| 11 | Hairs on outer surface of calyx (distribution): |
| | more or less absent (1); |
| | scattered in lower portion of calyx (2); |
| | scattered over the whole calyx (3); |
| | concentrated around the middle of the calyx (4); |
| | scattered over the whole calyx in bud but becoming more or less glabrous with age (5); or |
| | scattered over the abaxial half of the calyx and lobe (6) |
| 12 | Hairs on the calyx rim (presence): |
| | present only in the notch (1); |
| | present as tiny, scattered hairs towards apex of calyx lobes (2); or |
| | present on entire rim (3) |
| 13 | Hairs on the calyx rim (length): mm long |
| 14 | Hairs on calyx rim (density) (hairs in the notch are not included): |
| | absent (1); |
| | very sparse = some calyces on a specimen will have a few hairs on the calyx rim (2); |
| | sparse = need high power to find them, but present and spread out (3); or |
| | dense = hairs side by side (4) |
| 15 | Calyx (shape of lobes in bud): |
| | lobes clasping (1); |
| | lobes reflexed (3); or |
| | lobes neither clasping nor reflexed (2) |
| 16 | Calyx (tube length): mm long |
| 17 | Hairs on the corolla rim (density): |
| | absent (1); |
| | very sparse = obviously not side by side (2); |
| | sparse = side by side (3); or |
| | dense = hairs crowded together (4) |
| 18 | Hairs on the corolla rim (length): mm long |

[a] See Fig. S2

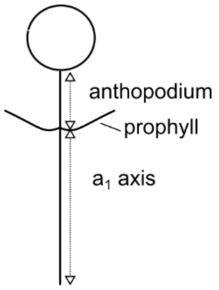


**Fig. S2** Structure of the ultimate inflorescence unit in *Prostanthera*, a uniflorescence *sensu* Conn (1984).

**Table S3 Morphological data used in PATN analysis**

The leading letter of the OTU (operational taxonomic unit) code is generally the leading letter of the specific epithet of the *Prostanthera* species expected at that site (C = *P. cineolifera*, H = *P.* sp. Hawkesbury, L = *P. lanceolata*, N = *Prostanthera* sp., O = *P. ovalifolia*, R = *P.* sp. Oxley Wild Rivers National Park, W = *P.* sp. Olney State Forest) except TB = Tabbimoble Creek. The specimens from TB had been determined as *P. cineolifera* but were considered by us to be unknown. The second and third letters are codes for the collection site. The digit is the number of the specimen from that site. Codes in bold indicate a type specimen. A dash indicates missing data. See Table S2 for characters. Herbarium codes follow Index Herbariorum (see http://sweetgum.nybg.org/ih/ accessed 14 December 2018.

| OTU | Herbarium accession | Character | | | | | | | | | | | | | | | | | |
|---|---|---|---|---|---|---|---|---|---|---|---|---|---|---|---|---|---|---|---|
| | | 1 | 2 | 3 | 4 | 5 | 6 | 7 | 8 | 9 | 10 | 11 | 12 | 13 | 14 | 15 | 16 | 17 | 18 |
| Cbb1 | n/a | 2 | 2 | 2.03 | 0.43 | 0.8 | 2 | | 3.67 | 2.25 | 3.25 | 1 | 3 | 0.1 | 4 | 1 | 2.25 | 1 | 0 |
| Cbn1 | NE 106411 | 2 | 2 | 3 | 0.95 | 0.8 | 2 | 1.3 | 1.9 | 1.6 | 2.9 | 1 | 3 | 0.14 | 4 | 1 | 2 | 1 | 0 |
| Ccc1 | NSW 424206 | 2 | 2 | 2.4 | 0.62 | 1.1 | 2 | | 3.75 | 2.83 | 5.5 | 6 | 3 | 0.1 | 4 | 1 | 2.3 | 1 | 0 |
| Cgr1 | NSW 227965 | 2 | 2 | 1.62 | 0.55 | 0.9 | 2 | | 2.33 | 1.67 | 3.25 | 1 | 3 | 0.2 | 4 | 1 | 2.33 | 1 | 0 |
| Cma1 | NSW 134412 | 2 | 2 | 1.6 | 0.59 | 0.65 | 2 | | 3.58 | 2.58 | 6.2 | 1 | 3 | 0.18 | 4 | | 2.33 | 1 | 0 |
| Cmc1 | NSW 134411 | 2 | 2 | 2.17 | 0.72 | 1 | 2 | | 3.67 | 2.58 | 6.3 | 1 | 3 | 0.24 | 4 | 1 | 2.42 | 1 | 0 |
| Csc1 | NSW 227969 | 2 | 2 | 1.98 | 0.55 | 0.9 | 1 | 1.67 | 2.33 | 1.67 | 3.25 | 5 | 3 | 0.14 | 4 | 1 | | 1 | 0 |
| **Csi-T** | NSW 134514 | 2 | 2 | 2.5 | 0.88 | 1.07 | 2 | | 3.42 | 2.25 | 8 | 1 | 3 | 0.3 | 4 | | 2.17 | 3 | 0.06 |
| Csw1 | NE 106008 | 2 | 2 | 2.4 | 0.75 | 1 | 2 | 1 | 3.4 | 2.7 | 4.2 | 1 | 3 | 0.12 | 4 | 1 | 4.25 | 1 | 0 |
| Cub1 | n/a | 2 | 2 | 2.75 | 0.92 | 1.03 | 2 | | 3.5 | 2.33 | 4.7 | 1 | 3 | 0.24 | 4 | 1 | 2.33 | 2 | 0.03 |
| Cwm1 | NE 106345 | 2 | 2 | 2.75 | 0.95 | 1.2 | 2 | | 2.5 | 1.8 | 2.25 | 6 | 3 | 0.14 | 4 | 1 | | 1 | 0 |
| Cwm2 | NE 106346 | 2 | 2 | 2.8 | 0.9 | 1.1 | 2 | 0.8 | 3.25 | 2.2 | 4.3 | 6 | 3 | 0.14 | 4 | 1 | 2.2 | 4 | 0 |
| Cwr1 | NE 105284 | 2 | 2 | 1.65 | 0.65 | 0.8 | 2 | | 3.8 | 2.7 | 3.2 | 6 | 3 | 0.12 | 4 | 1 | 2.4 | 3 | 0.04 |
| Hbc1 | NE 106340 | 1 | 2 | 2.55 | 0.55 | 1.1 | 1 | 2.75 | 1.2 | 1 | 1.8 | 4 | 3 | 0.05 | 4 | 2 | 1.75 | 3 | 0.05 |
| Hbc2 | NE 106339 | 1 | 2 | 2.33 | 0.52 | 1.07 | 1 | 2.92 | 1.58 | 1.17 | 3.5 | 3 | 3 | 0.04 | 4 | 2 | 2.25 | 4 | 0.1 |
| Hmo1 | NE 106006 | 1 | 2 | 2.93 | 0.78 | 1.1 | 1 | 2.5 | 0.92 | 0.67 | 2.83 | 3 | 3 | 0.04 | 4 | 2 | 2.08 | 4 | 0.06 |
| Hon1 | NE 106342 | 1 | 2 | 2.2 | 0.55 | 0.9 | 1 | 3 | 1 | 0.9 | 2.8 | 3 | 3 | 0.05 | 4 | 2 | 1.75 | 4 | 0.05 |
| Hon2 | NE 106212 | 1 | 2 | 1.85 | 0.55 | 0.8 | 1 | 2.75 | 0.8 | 0.7 | 2.1 | 4 | 3 | 0.07 | 4 | 2 | | 4 | 0.12 |
| Hon3 | NE 106385 | 1 | 2 | 2.93 | 0.7 | 1 | 1 | 3 | 1.92 | 1.67 | 4.25 | 3 | 3 | 0.04 | 4 | 2 | 2.17 | 4 | 0.08 |
| Lbm1 | NE 105873 | 1 | 1 | 2.4 | 1 | 1.3 | 2 | 1 | 2.25 | 1.8 | 3.8 | 1 | 1 | 0.04 | 2 | 3 | 2.25 | 3 | 0.02 |
| Lbm2 | NE 105874 | 1 | 1 | 2.15 | 0.8 | 1.1 | 2 | 0.75 | 2.1 | 1.7 | 4.3 | 1 | 1 | 0.05 | 2 | 3 | 2.6 | 3 | 0.05 |
| Lcr1 | NE 106857 | 1 | 1 | 1.57 | 0.63 | 0.6 | 2 | | 2 | 1.5 | 6.5 | 1 | 3 | 0.04 | 2 | 3 | 2.42 | 2 | 0.03 |
| Lma1 | NE 83318 | 1 | 1 | 1.22 | 0.43 | 0.38 | 2 | | 1.92 | 0.47 | 4 | 1 | 2 | 0.04 | 2 | 3 | 2.58 | 1 | 0 |

| OTU | Herbarium accession | Character | | | | | | | | | | | | | | | | | |
|---|---|---|---|---|---|---|---|---|---|---|---|---|---|---|---|---|---|---|---|
| | | **1** | **2** | **3** | **4** | **5** | **6** | **7** | **8** | **9** | **10** | **11** | **12** | **13** | **14** | **15** | **16** | **17** | **18** |
| Lmf1 | NE 105877 | 1 | 1 | 2.5 | 0.8 | 1.1 | 2 | 1.75 | 2.7 | 2 | 6.7 | 1 | 3 | 0.04 | 2 | 3 | 2.5 | 3 | 0.05 |
| Lmf2 | NE 105876 | 1 | 1 | 2.8 | 0.85 | 1.2 | 2 | 0.75 | 2.1 | 1.2 | 6.7 | 1 | 1 | 0 | 1 | 3 | 2.1 | 4 | 0.04 |
| Lny1 | NE 100319 | 1 | 1 | 1.82 | 0.63 | 0.9 | 2 | | 2.33 | 1.17 | 6.5 | 1 | 3 | 0.07 | 3 | 3 | 2.67 | 3 | 0.06 |
| Lsw1 | NE 106000 | 1 | 1 | 2.75 | 0.65 | 1.4 | 2 | | 1.8 | 1.2 | 6.3 | 1 | 3 | 0.04 | 2 | 3 | 2.4 | 3 | 0.05 |
| Ltm1 | NE 105880 | 1 | 1 | 2.5 | 0.8 | 1 | 2 | 1 | 1.75 | 1.25 | 6.5 | 1 | 1 | 0 | 1 | 3 | 2.4 | | 0.04 |
| Ltm2 | NE 105883 | 1 | 1 | 2.7 | 0.8 | 1.4 | 2 | | 2.6 | 1.4 | 9 | 1 | 1 | 0 | 1 | 3 | 2.5 | 3 | 0.04 |
| Lbs1 | BRI 313225 | 1 | 1 | 2.8 | 1.11 | 1.23 | 2 | | 1.5 | 1.17 | 6.5 | 1 | 3 | 0.09 | 4 | | 2.75 | 3 | 0.07 |
| Omm1 | NE 106180 | 3 | 1 | 1.7 | 0.58 | 0.75 | 2 | | 2.33 | 1.83 | 6.5 | 2 | 3 | 0.04 | 3 | 3 | 2.6 | 3 | 0.04 |
| Omm2 | NE 106182 | 3 | 1 | 1.85 | 0.8 | 0.7 | 2 | 1 | 1.75 | 1.42 | | 2 | 3 | 0.04 | 2 | 3 | 2.9 | 4 | 0.05 |
| Omm3 | NE 106181 | 3 | 2 | 1.5 | 0.62 | 0.67 | 2 | | 1.58 | 1.35 | 4.7 | 3 | 3 | 0.03 | 2 | 3 | 2.67 | 3 | 0.02 |
| Omn1 | NE 106207 | 1 | 2 | 1.6 | 0.5 | 0.65 | 2 | 1.2 | 2.2 | 1 | 4.3 | 1 | 3 | 0.05 | 2 | 3 | 2.7 | 3 | 0.04 |
| Omn2 | NE 106206 | 1 | 1 | 1.7 | 0.6 | 0.8 | 2 | 1.25 | 2.4 | 1.83 | 5.7 | 1 | 3 | 0.06 | 2 | 3 | 2.75 | 3 | 0.04 |
| Omn3 | NE 106203 | 1 | 1 | 2.25 | 0.65 | 0.9 | 2 | 1.9 | 1.9 | 1.5 | 2.4 | 1 | 3 | 0.05 | 2 | 3 | 2.8 | 3 | 0.05 |
| Omn4 | AQ 176156 | 1 | 1 | 1.05 | 0.43 | 0.43 | 2 | | 1.5 | 1.25 | 2.5 | 1 | 2 | 0.03 | 2 | 3 | 2.92 | 4 | 0.03 |
| Oms1 | NE 106213 | 1 | 2 | 1.5 | 0.5 | 0.7 | 2 | 0.6 | 1.6 | 1.1 | 2.3 | 1 | 3 | 0.02 | 2 | 3 | 2.7 | 4 | 0.04 |
| Oms2 | NE 106222 | 1 | 2 | 1.9 | 0.48 | 0.7 | 2 | 1.1 | 2.1 | 1.2 | | 1 | 3 | 0.05 | 2 | 3 | 2.9 | 3 | 0.02 |
| Oms3 | NE 106214 | 1 | 2 | 1.38 | 0.43 | 0.38 | 2 | | 1.83 | 1.25 | 3.5 | 1 | 1 | 0 | 1 | 3 | 3 | 4 | 0.06 |
| Oms4 | NE 106220 | 1 | 2 | 1.55 | 0.48 | 0.68 | 2 | | 2.83 | 1.25 | 5 | 1 | 1 | 0 | 2 | 3 | 2.75 | 4 | 0.06 |
| Rpr1 | NE 107073 | 1 | 2 | 1.55 | 0.63 | 0.55 | 1 | 2.5 | 0.75 | 0.5 | 1.83 | 3 | 3 | 0.09 | 4 | 2 | 2.58 | 4 | 0.35 |
| TB1 | NE 105986 | 1 | 1 | 1.85 | 0.5 | 1.75 | 2 | | 1.4 | 0.9 | 4.5 | 1 | 1 | 0.05 | 2 | 3 | 2 | 3 | 0.04 |
| Wmg1 | NE 49860 | 1 | 2 | 3.2 | 1.15 | 1 | 1 | 1.8 | 1 | 0.8 | 1.75 | 4 | 3 | 0.05 | 4 | 2 | 2 | 4 | 0.14 |
| Wol1 | NE 106323 | 1 | 2 | 3.6 | 1.2 | 1.1 | 1 | 1.8 | 0.5 | 0 | 1.6 | 1 | 3 | 0.04 | 3 | 2 | 1.4 | 4 | 0.04 |
| Wol2 | NE 106320 | 1 | 2 | 2.95 | 1 | 1.2 | 1 | 1.9 | 0.6 | 0.5 | 1.3 | 1 | 3 | 0.02 | 3 | 2 | 1.5 | 4 | 0.04 |
| Wol3 | NE 106220 | 1 | 2 | 1.85 | 0.83 | 0.68 | 1 | 2.08 | 0.8 | 0.7 | 2 | 1 | 3 | 0.08 | 3 | 2 | 1.58 | 4 | 0.08 |
| Wwf1 | CBG 003321 | 1 | 2 | 2.57 | 1.07 | 0.8 | 1 | 3.1 | 1.25 | 0.83 | 2 | 4 | 3 | 0.1 | 4 | 2 | 2.5 | 4 | 0.1 |

**Extra morphometric ordination plots – includes Figs S3 and S4**

The arrowed specimen in Fig. S3, BRI 313225, was collected by H. Streimann from Wilsons Creek Road, Mt Boss State Forest, New South Wales in October 1978 and is also arrowed in Fig. S4. This specimen had much wider leaves, a longer calyx tube and longer hairs on the calyx and corolla rims than is usual for the *Prostanthera ovalifolia/lanceolata* group. These features may be biogeographic variation. This specimen was not included in the molecular analysis of species boundaries. Further study of the *Prostanthera ovalifolia/lanceolata* group is needed to thoroughly explore possible species boundaries for this group.

BRI 313225 was not searchable on AVH (as of 9 March 2023). The duplicate, CBG 7809690.1, is searchable.

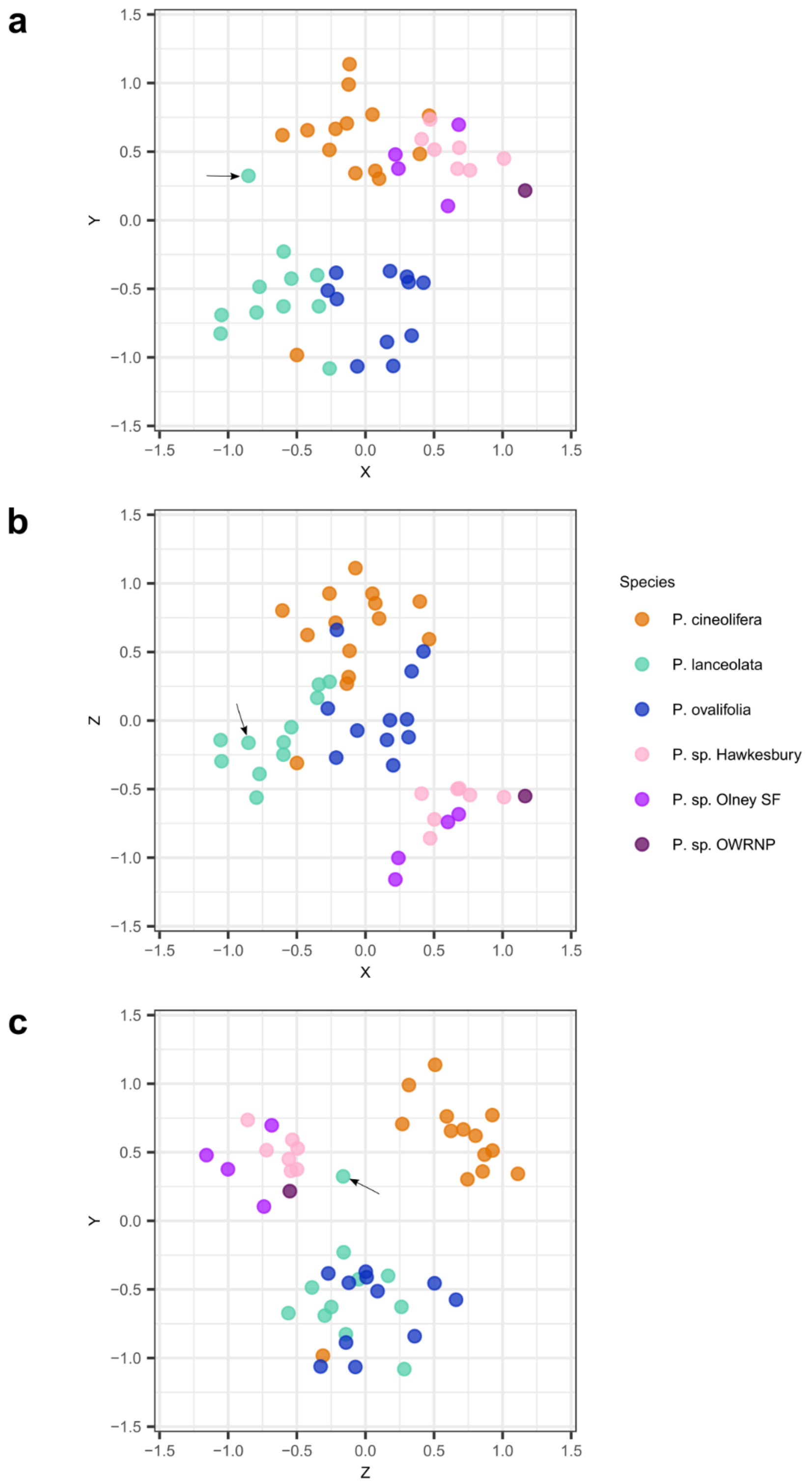


**Fig. S3** Morphometric ordination plot of semi-strong hybrid multidimensional scaling results. **a** Axis Y v. Axis Z, **b** Axis Z v. Axis X, **c** Axis Y v. Axis Z. The arrowed specimen, BRI 313225, is part of the cluster of the *Prostanthera ovalifolia/lanceolata* group. *P*. sp. Olney SF = *P*. sp. Olney State Forest, *P*. sp. OWRNP = *P*. sp. Oxley Wild Rivers National Park. Note: the arrowed specimen is not as remote from the *P. lanceolata* and *P. ovalifolia* group as it seems. See Fig. S4.

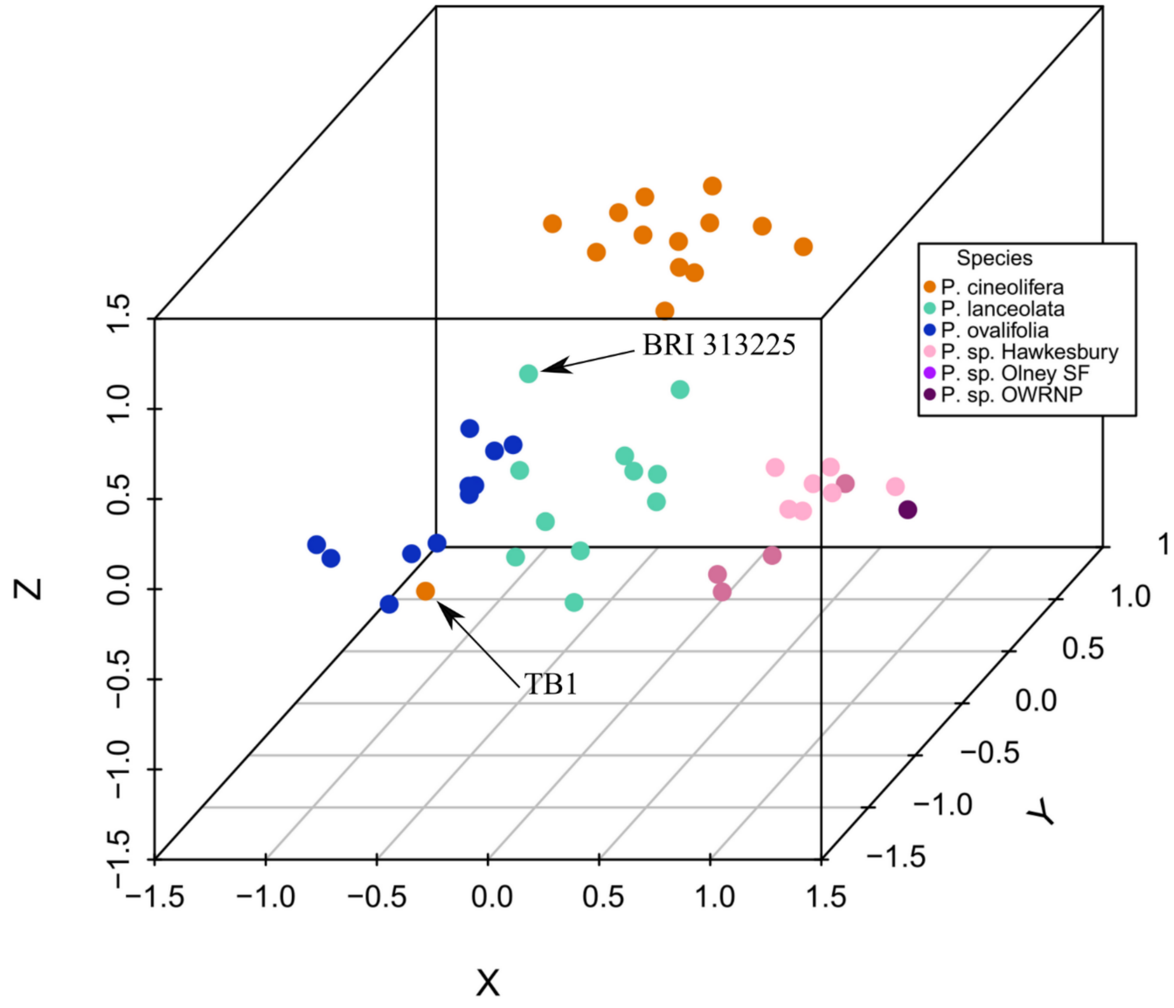


**Fig. S4** Three-dimensional morphometric ordination plot of semi-strong hybrid multidimensional scaling results. BRI 313225 is arrowed as is TB1 from Tabbimoble Creek. *P*. sp. Olney SF = *P*. sp. Olney State Forest, *P*. sp. OWRNP = *P*. sp. Oxley Wild Rivers National Park.

**Table S4 Details of samples sent to DArT**

The samples were randomised on the plates. For presentation here, the samples have been ordered in Clades, species and collection location. Three analyses were requested: Analysis 1 to include all samples, Analysis 2 to include only Clade J samples and Analysis 3 to include only Clade LAS samples. All samples were leaf tissue. In the Treatment column, Freeze dried indicates the leaf tissue was frozen and then freeze dried, Si Gel indicates the leaf tissue was dried in Silica Gel and Dried indicates the leaf tissue was from good quality herbarium specimens. The collector number plus Plate ID became the Individual ID used in the report, for example the specimen with collector number RLP152 on Plate 1 has an individual ID of RLP152-1 throughout the report. Ck = Creek, Mtn/s = Mountain/s, NP = National Park, NR = Nature Reserve, R = River, Rd = Road, SF = State Forest.

| Clade | Plate ID | Row | Column | Entity | Collector number | Treatment | Location |
|---|---|---|---|---|---|---|---|
| J | 1 | A | 10 | *Prostanthera cineolifera* | RLP152 | Freeze dried | Bees Nest Ridge_1 |
| J | 1 | C | 8 | *P. cineolifera* | RLP152a | Freeze dried | Bees Nest Ridge_1 |
| J | 1 | C | 9 | *P. cineolifera* | RLP152b | Freeze dried | Bees Nest Ridge_1 |
| J | 2 | D | 11 | *P. cineolifera* | RLP152c | Freeze dried | Bees Nest Ridge_1 |
| J | 2 | B | 5 | *P. cineolifera* | RLP153 | Freeze dried | Bees Nest Ridge_1 |
| J | 2 | B | 3 | *P. cineolifera* | RLP154 | Freeze dried | Bees Nest Ridge_1 |
| J | 1 | D | 3 | *P. cineolifera* | RLP155 | Freeze dried | Bees Nest Ridge_2 |
| J | 1 | H | 4 | *P. cineolifera* | RLP155a | Freeze dried | Bees Nest Ridge_2 |
| J | 2 | G | 10 | *P. cineolifera* | RLP155c | Freeze dried | Bees Nest Ridge_2 |
| J | 2 | E | 5 | *P. cineolifera* | RLP156 | Freeze dried | Bees Nest Ridge_2 |
| J | 2 | H | 10 | *P. cineolifera* | RLP158 | Freeze dried | Bees Nest Ridge_3 |
| J | 2 | A | 11 | *P. cineolifera* | RLP158a | Freeze dried | Bees Nest Ridge_3 |
| J | 2 | C | 7 | *P. cineolifera* | RLP158b | Freeze dried | Bees Nest Ridge_3 |
| J | 2 | E | 8 | *P. cineolifera* | RLP158c | Freeze dried | Bees Nest Ridge_3 |
| J | 2 | F | 8 | *P. cineolifera* | RLP158d | Freeze dried | Bees Nest Ridge_3 |
| J | 2 | A | 10 | *P. cineolifera* | RLP158e | Freeze dried | Bees Nest Ridge_3 |
| J | 1 | C | 2 | *P. cineolifera* | RLP160 | Freeze dried | Bees Nest Ridge_4 |
| J | 1 | H | 10 | *P. cineolifera* | RLP160a | Freeze dried | Bees Nest Ridge_4 |
| J | 1 | A | 8 | *P. cineolifera* | RLP160b | Freeze dried | Bees Nest Ridge_4 |
| J | 1 | G | 2 | *P. cineolifera* | RLP160c | Freeze dried | Bees Nest Ridge_4 |
| J | 2 | A | 9 | *P. cineolifera* | RLP160d | Freeze dried | Bees Nest Ridge_4 |
| J | 2 | B | 2 | *P. cineolifera* | RLP160e | Freeze dried | Bees Nest Ridge_4 |

| Clade | Plate ID | Row | Column | Entity | Collector number | Treatment | Location |
|---|---|---|---|---|---|---|---|
| J | 1 | H | 11 | *P. cineolifera* | RLP160f | Freeze dried | Bees Nest Ridge_4 |
| J | 2 | E | 11 | *P. cineolifera* | RLP148 | Freeze dried | Broke Rd |
| J | 1 | H | 7 | *P. cineolifera* | RLP142 | Si Gel | Broken Back Trail |
| J | 1 | A | 9 | *P. cineolifera* | RLP143 | Si Gel | Broken Back Trail |
| J | 1 | H | 2 | *P. cineolifera* | RLP144 | Si Gel | Broken Back Trail |
| J | 1 | D | 2 | *P. cineolifera* | RLP145 | Si Gel | Broken Back Trail |
| J | 2 | B | 12 | *P. cineolifera* | RLP146 | Si Gel | Broken Back Trail |
| J | 1 | G | 10 | *P. cineolifera* | RLP147 | Freeze dried | Broken Back Trail |
| J | 1 | B | 2 | *P. cineolifera* | RLP228 | Freeze dried | Cousins Creek |
| J | 1 | B | 4 | *P. cineolifera* | RLP228A | Freeze dried | Cousins Creek |
| J | 1 | H | 8 | *P. cineolifera* | RLP228B | Freeze dried | Cousins Creek |
| J | 1 | G | 7 | *P. cineolifera* | RLP228C | Freeze dried | Cousins Creek |
| J | 1 | B | 5 | *P. cineolifera* | RLP228D | Freeze dried | Cousins Creek |
| J | 2 | C | 8 | *P. cineolifera* | RLP228E | Freeze dried | Cousins Creek |
| J | 1 | E | 8 | *P. cineolifera* | RLP229 | Freeze dried | Cousins Creek |
| J | 1 | D | 8 | *P. cineolifera* | RLP229A | Freeze dried | Cousins Creek |
| J | 1 | F | 2 | *P. cineolifera* | RLP135 | Si Gel | Sawpit Rd |
| J | 1 | H | 6 | *P. cineolifera* | RLP136 | Si Gel | Sawpit Rd |
| J | 1 | E | 10 | *P. cineolifera* | RLP137 | Si Gel | Sawpit Rd |
| J | 1 | B | 8 | *P. cineolifera* | RLP138 | Si Gel | Sawpit Rd |
| J | 1 | G | 5 | *P. cineolifera* | RLP139 | Si Gel | Sawpit Rd |
| J | 2 | D | 10 | *P. cineolifera* | RLP140 | Si Gel | Sawpit Rd |
| J | 1 | G | 8 | *P. cineolifera* | RLP207 | Freeze dried | Wallaby Rocks |
| J | 2 | A | 8 | *P. cineolifera* | RLP207 | Freeze dried | Wallaby Rocks |
| J | 1 | C | 4 | *P. cineolifera* | RLP208 | Freeze dried | Wallaby Rocks |
| J | 1 | C | 6 | *P. cineolifera* | RLP210 | Freeze dried | Wallaby Rocks |
| J | 1 | E | 3 | *P. cineolifera* | RLP177 | Freeze dried | Wingen Maid |
| J | 1 | G | 11 | *P. cineolifera* | RLP178 | Freeze dried | Wingen Maid |
| J | 1 | H | 9 | *P. cineolifera* | RLP179 | Freeze dried | Wingen Maid |
| J | 1 | E | 2 | *P. cineolifera* | RLP180 | Freeze dried | Wingen Maid |

| Clade | Plate ID | Row | Column | Entity | Collector number | Treatment | Location |
|---|---|---|---|---|---|---|---|
| J | 1 | E | 6 | *P. cineolifera* | RLP181 | Freeze dried | Wingen Maid |
| J | 1 | F | 8 | *P. cineolifera* | RLP182 | Freeze dried | Wingen Maid |
| J | 2 | E | 1 | *P. cineolifera* | RLP182 | Freeze dried | Wingen Maid |
| J | 2 | C | 4 | *P. cotinifolia* A.Cunn. ex Benth. | NJS543 | Dried | Warrumbungles |
| J | 1 | H | 1 | *P. incisa* | NJS274 | Dried | Blue Mtns |
| J | 2 | E | 6 | *P. incisa* | NJS274 | Dried | Blue Mtns |
| J | 1 | D | 4 | *P. incisa* | JJB3532 | Freeze dried | Werrikimbe NP |
| J | 2 | A | 2 | *P. incisa* | JJB3532 | Freeze dried | Werrikimbe NP |
| J | 1 | D | 12 | *P. incisa* | JJB3532a | Freeze dried | Werrikimbe NP |
| J | 2 | C | 9 | *P. incisa* | JJB3532a | Freeze dried | Werrikimbe NP |
| J | 2 | F | 2 | *P. incisa* | JJB3532e | Freeze dried | Werrikimbe NP |
| J | 1 | F | 12 | *P. lanceolata* | RLP110 | Freeze dried | Bar Mountain |
| J | 2 | F | 1 | *P. lanceolata* | RLP110 | Freeze dried | Bar Mountain |
| J | 1 | D | 9 | *P. lanceolata* | RLP110a | Freeze dried | Bar Mountain |
| J | 1 | H | 5 | *P. lanceolata* | PGW1890 | Dried | Carrai |
| J | 1 | E | 1 | *P. lanceolata* | JJB3542 | Freeze dried | Middle Brother Mtn NP |
| J | 2 | H | 9 | *P. lanceolata* | JJB3542 | Freeze dried | Middle Brother Mtn NP |
| J | 1 | A | 12 | *P. lanceolata* | JJB3542a | Freeze dried | Middle Brother Mtn NP |
| J | 1 | B | 7 | *P. lanceolata* | RLP113 | Freeze dried | Minyon Falls |
| J | 2 | G | 8 | *P. lanceolata* | RLP113 | Freeze dried | Minyon Falls |
| J | 1 | G | 4 | *P. lanceolata* | RLP114 | Freeze dried | Minyon Falls |
| J | 1 | A | 6 | *P. lanceolata* | JJB3563 | Freeze dried | New England NP |
| J | 2 | G | 2 | *P. lanceolata* | JJB3563 | Freeze dried | New England NP |
| J | 1 | B | 1 | *P. lanceolata* | JJB3563a | Freeze dried | New England NP |
| J | 1 | A | 7 | *P. lanceolata* | NJS239 | Dried | Nymboida |
| J | 2 | E | 7 | *P. lanceolata* | RLP121 | Freeze dried | Pillar Valley |
| J | 1 | F | 7 | *P. lanceolata* | RLP121a | Freeze dried | Pillar Valley |
| J | 1 | C | 10 | *P. lanceolata* | RLP121e | Freeze dried | Pillar Valley |
| J | 1 | F | 4 | *P. lanceolata* | RLP122 | Freeze dried | Sherwood NR |
| J | 2 | F | 7 | *P. lanceolata* | RLP122 | Freeze dried | Sherwood NR |

| Clade | Plate ID | Row | Column | Entity | Collector number | Treatment | Location |
|---|---|---|---|---|---|---|---|
| J | 1 | E | 12 | *P. lanceolata* | RLP122a | Freeze dried | Sherwood NR |
| J | 1 | H | 3 | *P. lanceolata* | RLP104 | Freeze dried | Tamborine |
| J | 1 | E | 7 | *P. lanceolata* | RLP107 | Freeze dried | Tamborine |
| J | 2 | F | 9 | *P. lanceolata* | RLP107 | Freeze dried | Tamborine |
| J | 2 | H | 1 | *P. latifolia* (Benth.) Domin | IRT13512 | Dried | Tapin Tops |
| J | 2 | H | 8 | *P. latifolia* | NJS424 | Dried | Tapin Tops |
| J | 1 | F | 11 | *P. ovalifolia* | RLP188 | Freeze dried | Mt Maria |
| J | 1 | B | 3 | *P. ovalifolia* | RLP188a | Freeze dried | Mt Maria |
| J | 1 | G | 1 | *P. ovalifolia* | RLP188b | Freeze dried | Mt Maria |
| J | 1 | F | 9 | *P. ovalifolia* | RLP188c | Freeze dried | Mt Maria |
| J | 1 | C | 12 | *P. ovalifolia* | RLP188d | Freeze dried | Mt Maria |
| J | 2 | D | 6 | *P. ovalifolia* | RLP188e | Freeze dried | Mt Maria |
| J | 2 | G | 9 | *P. ovalifolia* | RLP188f | Freeze dried | Mt Maria |
| J | 1 | E | 5 | *P. ovalifolia* | RLP184 | Freeze dried | Mt Ninderry |
| J | 1 | F | 5 | *P. ovalifolia* | RLP184a | Freeze dried | Mt Ninderry |
| J | 1 | E | 11 | *P. ovalifolia* | RLP184b | Freeze dried | Mt Ninderry |
| J | 1 | C | 5 | *P. ovalifolia* | RLP184c | Freeze dried | Mt Ninderry |
| J | 1 | B | 12 | *P. ovalifolia* | RLP184d | Freeze dried | Mt Ninderry |
| J | 2 | C | 3 | *P. ovalifolia* | RLP184e | Freeze dried | Mt Ninderry |
| J | 2 | E | 4 | *P. ovalifolia* | RLP185 | Freeze dried | Mt Ninderry |
| J | 1 | A | 5 | *P. ovalifolia* | RLP195 | Freeze dried | Mt Stanley |
| J | 1 | F | 10 | *P. ovalifolia* | RLP195a | Freeze dried | Mt Stanley |
| J | 2 | G | 3 | *P. ovalifolia* | RLP195b | Freeze dried | Mt Stanley |
| J | 1 | A | 3 | *P. ovalifolia* | RLP200 | Freeze dried | Mt Stanley |
| J | 1 | F | 1 | *P. ovalifolia* | RLP200a | Freeze dried | Mt Stanley |
| J | 2 | G | 5 | *P. ovalifolia* | RLP200b | Freeze dried | Mt Stanley |
| J | 1 | C | 1 | *P. ovalifolia* | RLP201 | Freeze dried | Mt Stanley |
| J | 1 | E | 4 | *P. ovalifolia* | RLP201a | Freeze dried | Mt Stanley |
| J | 2 | D | 12 | *P. ovalifolia* | RLP201b | Freeze dried | Mt Stanley |
| J | 2 | G | 7 | *P. petraea* B.J.Conn | JJB3540a | Si gel | Girraween |

| Clade | Plate ID | Row | Column | Entity | Collector number | Treatment | Location |
|---|---|---|---|---|---|---|---|
| J | 2 | B | 7 | *P. petraea* | JJB3540 | Si gel | Girraween |
| J | 2 | D | 4 | *P. rotundifolia* | JRN158 | Freeze dried | Genoa Falls |
| J | 2 | B | 8 | *P. rotundifolia* | JRN155 | Si gel | Launceston |
| J | 2 | B | 4 | *P. rotundifolia* | JRN155a | Dried | Launceston |
| J | 2 | C | 1 | *P*. sp. Barren Mtn | NJS366 | Dried | Barren Mtn |
| J | 2 | E | 3 | *P*. sp. Barren Mtn | NJS427 | Dried | Barren Mtn |
| J | 2 | C | 10 | *P*. sp. Barren Mtn | NJS428 | Dried | Barren Mtn |
| J | 2 | F | 6 | *P*. sp. Blue Mtns | JRN137 | Dried | Blue Mtns |
| J | 2 | D | 5 | *P*. sp. Blue Mtns | NJS542 | Dried | Blue Mtns |
| J | 1 | B | 9 | *P*. sp. Dandahra Ck | JJB3515 | Dried | Dandahra Ck |
| J | 2 | F | 4 | *P*. sp. Dandahra Ck | JJB3515 | Dried | Dandahra Ck |
| J | 1 | C | 7 | *P*. sp. Dandahra Ck | NJS296 | Dried | Dandahra Ck |
| J | 2 | F | 11 | *P*. sp. Dandahra Ck | NJS296 | Dried | Dandahra Ck |
| J | 2 | E | 9 | *P*. sp. Grampians | JRN104 | Dried | Grampians |
| J | 1 | C | 3 | *P*. sp. Hawkesbury | RLP167 | Freeze dried | Bicentenary Rd |
| J | 1 | D | 6 | *P*. sp. Hawkesbury | RLP168 | Freeze dried | Bicentenary Rd |
| J | 1 | F | 6 | *P*. sp. Hawkesbury | RLP169 | Freeze dried | Bicentenary Rd |
| J | 1 | D | 11 | *P*. sp. Hawkesbury | RLP170 | Freeze dried | Bicentenary Rd |
| J | 2 | A | 7 | *P*. sp. Hawkesbury | RLP171 | Freeze dried | Bicentenary Rd |
| J | 2 | A | 12 | *P*. sp. Hawkesbury | RLP172 | Freeze dried | Bicentenary Rd |
| J | 1 | A | 2 | *P*. sp. Hawkesbury | RLP206 | Freeze dried | Bar Point |
| J | 2 | C | 2 | *P*. sp. Hawkesbury | RLP206 | Freeze dried | Bar Point |
| J | 1 | D | 7 | *P*. sp. Hawkesbury | RLP206a | Freeze dried | Bar Point |
| J | 1 | F | 3 | *P*. sp. Hawkesbury | MRD105 | Si Gel | Old Northern Rd |
| J | 1 | A | 4 | *P*. sp. Hawkesbury | RLP173 | Si Gel | Old Northern Rd |
| J | 2 | D | 7 | *P*. sp. Hawkesbury | RLP173 | Si Gel | Old Northern Rd |
| J | 1 | E | 9 | *P*. sp. Hawkesbury | RLP173a | Freeze dried | Old Northern Rd |
| J | 1 | G | 6 | *P*. sp. Hawkesbury | RLP174 | Freeze dried | Old Northern Rd |
| J | 1 | G | 9 | *P*. sp. Hawkesbury | RLP174b | Freeze dried | Old Northern Rd |
| J | 2 | F | 3 | *P*. sp. Hawkesbury | RLP176 | Freeze dried | Old Northern Rd |

| Clade | Plate ID | Row | Column | Entity | Collector number | Treatment | Location |
|---|---|---|---|---|---|---|---|
| J | 2 | H | 11 | *P*. sp. Mt Marsh | LMC3770 | Dried | Mt Marsh |
| J | 1 | D | 5 | *P*. sp. Olney State Forest | RLP161 | Freeze dried | Walkers Ridge |
| J | 2 | D | 1 | *P*. sp. Olney State Forest | RLP161 | Freeze dried | Walkers Ridge |
| J | 1 | A | 11 | *P*. sp. Olney State Forest | RLP162 | Freeze dried | Walkers Ridge |
| J | 2 | D | 3 | *P*. sp. Olney State Forest | RLP162 | Freeze dried | Walkers Ridge |
| J | 1 | C | 11 | *P*. sp. Olney State Forest | RLP163 | Freeze dried | Walkers Ridge |
| J | 1 | A | 1 | *P*. sp. Olney State Forest | RLP164 | Freeze dried | Walkers Ridge |
| J | 1 | B | 6 | *P*. sp. Olney State Forest | RLP165 | Freeze dried | Walkers Ridge |
| J | 2 | E | 2 | *P*. sp. Olney State Forest | RLP166 | Freeze dried | Walkers Ridge |
| J | 2 | F | 10 | *P*. sp. Oxley Wild Rivers National Park | LMC4355 | Dried | Garabaldi Homestead |
| J | 2 | B | 1 | *P*. sp. Oxley Wild Rivers National Park | MFD4027 | Dried | Paradise Rocks |
| J | 2 | C | 5 | *P*. sp. Ulan | NJS292 | Dried | Ulan |
| J | 1 | B | 11 | Unknown | RLP119 | Freeze dried | Tabbimoble Ck |
| J | 2 | D | 9 | Unknown | RLP119 | Freeze dried | Tabbimoble Ck |
| J | 1 | D | 10 | Unknown | RLP120 | Freeze dried | Tabbimoble Ck |
| LAS | 2 | B | 9 | *P. lasianthos Labill. var. lasianthos* | DWL894 | Dried | Mt Wellington |
| LAS | 2 | H | 5 | *P. lasianthos Labill. var. lasianthos* | DWL895 | Dried | Mt Wellington |
| LAS | 2 | C | 12 | *P. lasianthos var. subcoriacea F.Muell. ex Benth.* | JRN 116 | Dried | Grampians |
| LAS | 2 | F | 12 | *P. lasianthos var. subcoriacea* | JRN115 | Dried | Grampians |
| LAS | 2 | A | 5 | *P*. sp. Australian Alps | JJB3371 | Si gel | Alpine Way |
| LAS | 2 | E | 12 | *P*. sp. Australian Alps | JJB3366 | Dried | Snowyridge Rd |
| LAS | 2 | A | 4 | *P*. sp. Australian Alps | JJB2564 | Dried | Thredbo R |
| LAS | 2 | G | 6 | *P. sp. Bald Mountain (M.S. Clemens AQ336575)* | IRT13128 | Dried | Donnybrook |
| LAS | 2 | B | 6 | *P. sp. Bald Mountain (M.S. Clemens AQ336575)* | IRT13129 | Dried | Donnybrook |
| LAS | 2 | H | 7 | *P. sp. Bald Mountain (M.S. Clemens AQ336575)* | IRT13130 | Dried | Donnybrook |
| LAS | 2 | C | 11 | *P. sp. Mount Kaputar (R.H.Cambage 5/11/1909)* | RLP232 | Freeze dried | Lindsay Rock Tops |
| LAS | 2 | C | 6 | *P. sp. Mount Kaputar (R.H.Cambage 5/11/1909)* | RLP232a | Freeze dried | Lindsay Rock Tops |
| LAS | 2 | H | 6 | *P. sp. Mount Kaputar (R.H.Cambage 5/11/1909)* | RLP232b | Freeze dried | Lindsay Rock Tops |
| LAS | 2 | G | 11 | *P*. sp. New England | JJB3579 | Freeze dried | Banksia Point |
| LAS | 2 | G | 1 | *P*. sp. New England | JJB3579a | Freeze dried | Banksia Point |

| Clade | Plate ID | Row | Column | Entity | Collector number | Treatment | Location |
|---|---|---|---|---|---|---|---|
| LAS | 2 | A | 6 | *P.* sp. New England | JJB3579b | Freeze dried | Banksia Point |
| LAS | 2 | D | 2 | *P.* sp. New England | JJB3580 | Freeze dried | Toms Cabin |
| LAS | 2 | A | 3 | *P.* sp. New England | JJB3580b | Freeze dried | Toms Cabin |
| LAS | 2 | H | 3 | *P. sp. Schofields Gap (I.R.Telford 12807)* | JJB3578 | Freeze dried | Cathedral Rock NP |
| LAS | 2 | D | 8 | *P. sp. Schofields Gap (I.R.Telford 12807)* | JJB3578b | Freeze dried | Cathedral Rock NP |
| LAS | 1 | G | 3 | *P. sp. Schofields Gap (I.R.Telford 12807)* | JJB3578c | Freeze dried | Cathedral Rock NP |
| LAS | 2 | B | 11 | *P. sp. Schofields Gap (I.R.Telford 12807)* | JJB3577 | Freeze dried | Deervale Loop Rd |
| LAS | 2 | H | 4 | *P. sp. Schofields Gap (I.R.Telford 12807)* | JJB3577a | Freeze dried | Deervale Loop Rd |
| LAS | 2 | B | 10 | *P. sp. Schofields Gap (I.R.Telford 12807)* | JJB3577b | Freeze dried | Deervale Loop Rd |
| LAS | 1 | D | 1 | *P. sp. Schofields Gap (I.R.Telford 12807)* | JJB3577c | Freeze dried | Deervale Loop Rd |
| LAS | 2 | G | 4 | *P. sp. Wollomombi Falls (J.B.Williams NE34800)* | JJB3582 | Freeze dried | Apsley Falls |
| LAS | 2 | A | 1 | *P. sp. Wollomombi Falls (J.B.Williams NE34800)* | JJB3582b | Freeze dried | Apsley Falls |
| LAS | 2 | H | 2 | *P. sp. Wollomombi Falls (J.B.Williams NE34800)* | JJB3581 | Freeze dried | Edgars Lookout |
| LAS | 2 | F | 5 | *P. sp. Wollomombi Falls (J.B.Williams NE34800)* | JJB3581a | Freeze dried | Edgars Lookout |
| LAS | 2 | E | 10 | *P. sp. Wollomombi Falls (J.B.Williams NE34800)* | JJB3581b | Freeze dried | Edgars Lookout |
| LAS | 1 | B | 10 | *P. sp. Wollomombi Falls (J.B.Williams NE34800)* | JJB3581c | Freeze dried | Edgars Lookout |

##### Technical duplicates including Table S5

The technical duplicates were included to test the reproducibility of the marker data in the DArTseq procedure. Eighteen of the samples on Plate 1 had a technical duplicate in Plate 2 (**Table S2**). Sixteen pairs of duplicates remaining after filtering. To examine the variability visually, the 'gl.pcoa' function in *dartR* (Gruber *et al.* 2017) was used to conduct PCA on the 16 pairs of technical duplicates (Fig. S5). PCA Axis 1 explained 25.0% of the total variance. Axis 2 explained 14.2% of the total variance. Axes 1-3 combined explained 51.4% of the total variance. Scrutiny of the PCA plots indicated that most duplicate pairs plotted together or very closely. Two pairs, RLP207 and JJB3515 (arrowed in Fig. 2-27), were separated by a small distance in the plots. This distance is an indication of possible errors or random missing data in the SNP data. This level of noise is trivial compared to the strength of clustering. There is no cause for concern in combining data from Plate 1 and data from Plate 2. Although results from the two plates are not identical, there are only small differences.

**Table S2** These individuals were represented on both plates. Ck = Creek, NP = National Park, Qld = Queensland, SF = State Forest, Rd = Road. All locations in New South Wales unless otherwise stated.

| Species | Voucher | Location |
|---|---|---|
| *Prostanthera cineolifera* | RLP182 | Wingen Maid |
| *P. cineolifera* | RLP207 | Wallaby Rocks |
| *P. lanceolata* | JJB3542 | Middle Brother Mtn NP |
| *P. lanceolata* | JJB3563 | New England NP |
| *P. lanceolata* | RLP107 | Tamborine Mtn, Qld |
| *P. lanceolata* | RLP110 | Bar Mountain |
| *P. lanceolata* | RLP113 | Minyon Falls |
| *P. lanceolata* | RLP119 | Tabbimoble Ck |
| *P. lanceolata* | RLP122 | Sherwood NR |
| *P. sp. Dandahra Ck* | JJB3515 | Dandahra Ck |
| *P. sp. Dandahra Ck* | NJS296 # | Dandahra Ck |
| *P.* sp. Hawkesbury | RLP173 # | Old Northern Rd |
| *P.* sp. Hawkesbury | RLP206 | Bar Point |
| *P.* sp. Olney State Forest | RLP161 | Walkers Ridge |
| *P.* sp. Olney State Forest | RLP162 | Walkers Ridge |
| *P. incisa* | JJB3532 | Werrikimbe NP |
| *P. incisa* | JJB3532a | Werrikimbe NP |
| *P. incisa* | NJS274 | Blue Mtns |

### One of the duplicate pair was filtered out.

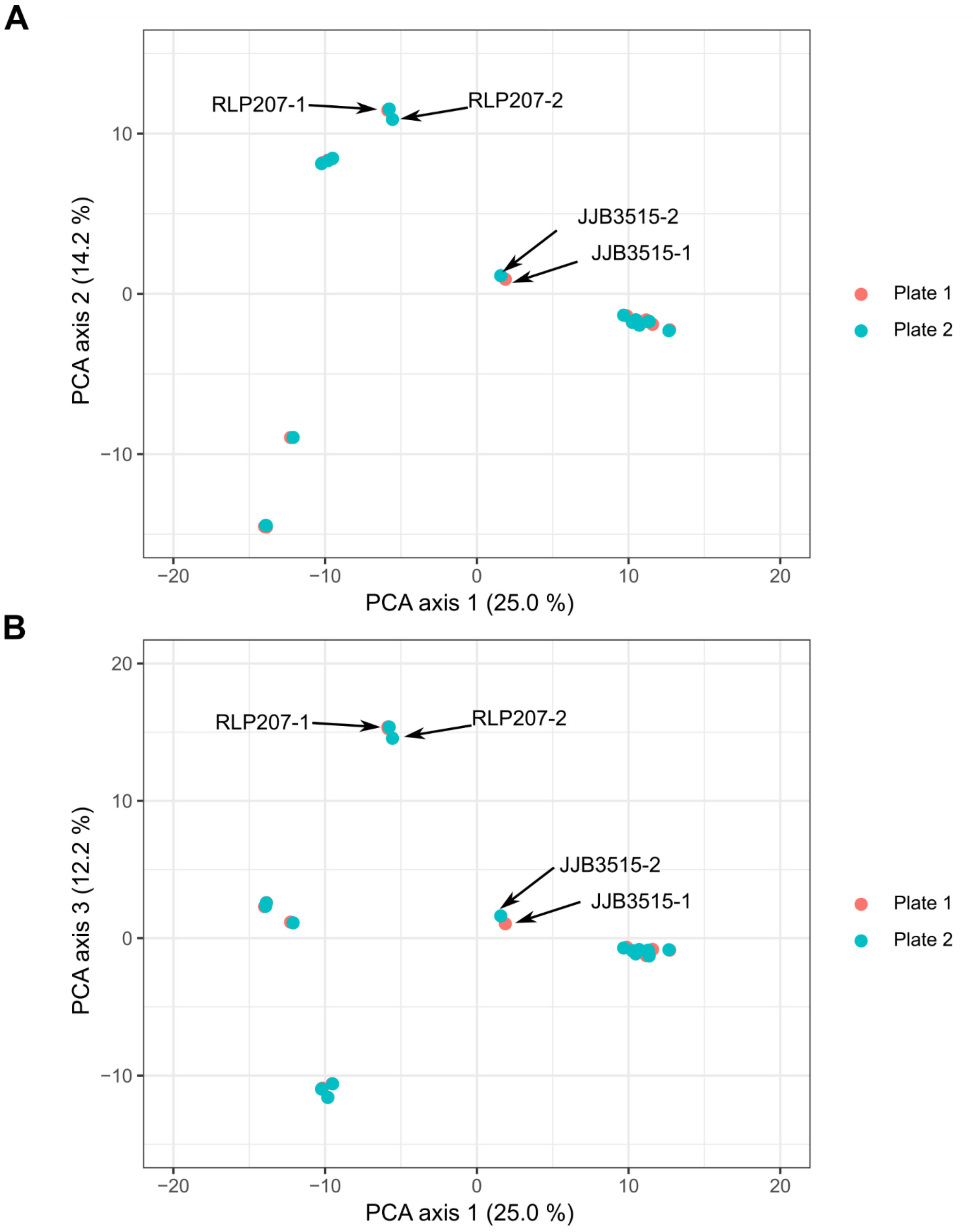


**Fig. S5** Principal Coordinates Analysis (PCA) of SNP data from the technical duplicates (6,198 loci from 32 individuals). **A** Axis 2 vs axis 1. **B** Axis 3 vs axis 1. The two duplicate pairs with the greatest separation are RLP207 and JJB3515 (arrowed). Note: RLP207-1 is barely visible as an orange sliver under a blue dot that represents RLP182-2. Percentage of genetic distance explained by each of the PCA axes is provided in parentheses.

**Anomalous specimens**

The complete dataset of Clade J samples with 12,123 loci across 148 individuals, was used for preliminary exploratory analysis. Molecular Principal Coordinates Analysis (PCA), a method of visualizing genetic similarity of individuals and populations (Gower 1971) was conducted on the complete dataset using the 'gl.pcoa' function in *dartR* (Gruber *et al.* 2017). The resulting plot (Fig. S) indicated that four specimens were anomalous, likely through errors in plate packing, considering their plate positions. As a result, the following samples were omitted:

- PGW1890-1 (Fig. S), expected to be labelled as *P. lanceolata*, plotted halfway between the *P. lanceolata* and *P. ovalifolia* cluster and the *P. cineolifera* cluster—strong indications that it was a mixed sample.
- RLP184-1 (Fig. S), expected to be *P. ovalifolia* from Mt Ninderry in southeastern Queensland, plotted halfway between the *P. lanceolata* and *P. ovalifolia* cluster and the *P. cineolifera* cluster—strong indications that it was a mixed sample.
- RLP195a-1 (Fig. S), expected to be *P. ovalifolia* from Mt Stanley in central Queensland, plotted partway towards the *P. cineolifera* cluster, indicating contamination of the RLP195a-1 sample with material from *P. cineolifera.*
- RLP206a-1 (Fig. S), expected to be *P*. sp. Hawkesbury from Bar Point (Lower Hawkesbury River), clustered with specimens of *P. ovalifolia* from Mt Maria and Mt Stanley in Central Queensland. These two species are morphologically very distinct. RLP206-1, collected from the same population as RLP206a-1, clustered with all the other individuals of *P*. sp. Hawkesbury, indicating that the sample was mislabeled.

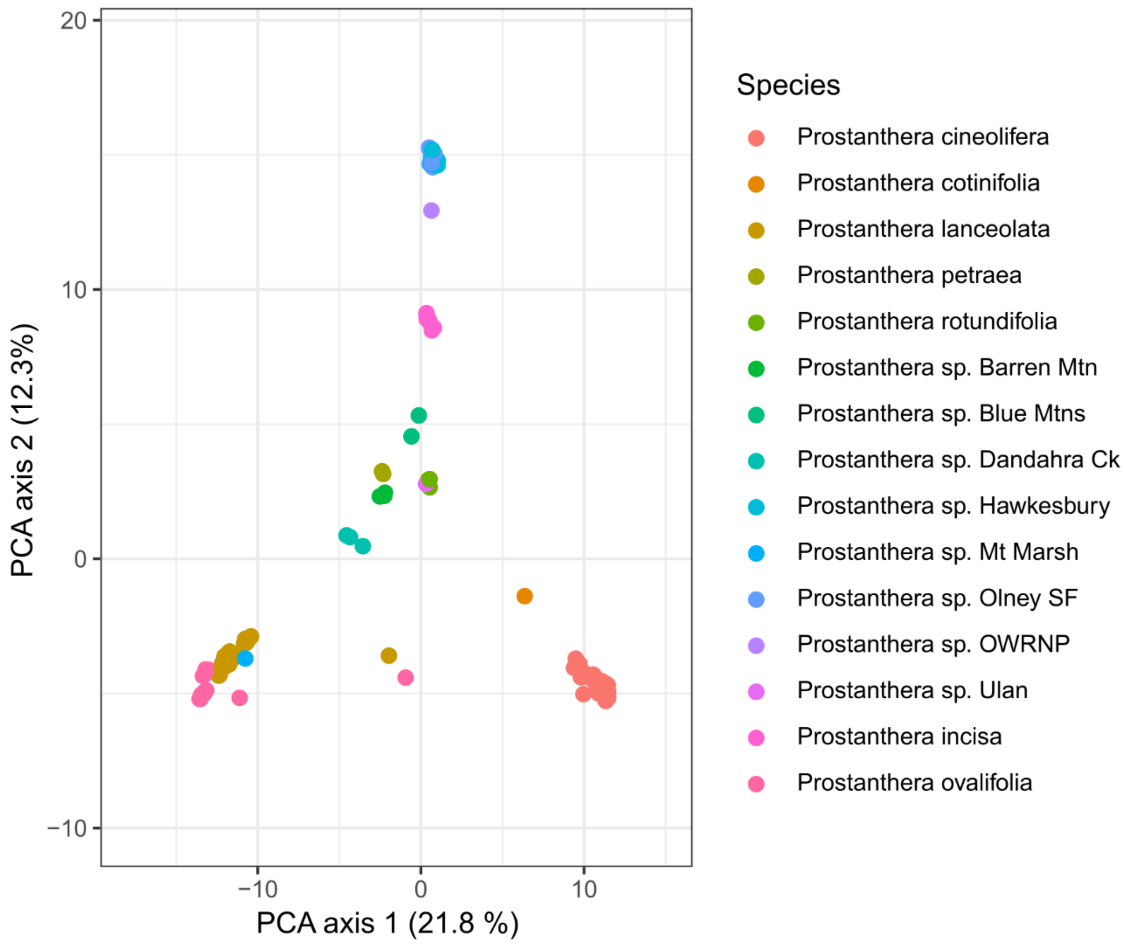


**Fig. S6** Principal Coordinates Analysis (PCA) of SNP data from the complete dataset (12,123 loci from 148 individuals). Arrowed individuals plotted in unexpected places: PGW1890-1 (labelled as *Prostanthera lanceolata*) and RLP184-1(labelled as *P. ovalifolia*) but clustering half way between the *P. lanceolata* and *P. ovalifolia* cluster and the *P. cineolifera* cluster; RLP195a-1 (labelled as *P. ovalifolia*) clustering partway towards the *P. cineolifera* cluster); and RLP206a-1 (labelled as *P.* sp. Hawkesbury), but clustering with specimens of *P. ovalifolia*—Note: RLP206a-1 is hidden under other individuals. *P.* sp. Olney SF = *P.* sp. Olney State Forest, *P.* sp. OWRNP = *P.* sp. Oxley Wild Rivers National Park.

#### Table S6 Filters used and the resulting numbers of individuals and loci

Where filters did not exclude uninformative monomorphic loci, this filter was applied separately.

| Filter name (threshold) | Number of individuals after filtering | Number of loci after filtering |
|---|---|---|
| Initial counts | 157 | 185,761 |
| Filter loci on repeatability (100%) | 157 | 174,078 |
| Filter loci on call rate (95%) | 157 | 17,095 |
| Filter individuals on call rate (90%) | 148 | 16,877 |
| Filter out loci with similar trimmed sequence tags (25%) | 148 | 12,123 |

#### Table S7 Samples removed during the filtering process due to low call rates

Ck = Creek, NP = National Park, Rd = Road. The suffix to the individual ID (1 or 2) indicates the plate number.

| **Individual ID-plate number** | **Collection site** | **Treatment** | **Species** |
|---|---|---|---|
| RLP210-1 | Wallaby Rocks | Freeze dried | *Prostanthera cineolifera* |
| NJS296-1 | Dandahra Ck | Dried | *P.* sp. Dandahra Ck |
| MFD4027-2 | Paradise Rocks | Dried | *P.* sp. Oxley Wild Rivers National Park |
| RLP158b-2 | Bees Nest Ridge | Freeze dried | *P. cineolifera* |
| RLP152c-2 | Bees Nest Ridge | Freeze dried | *P. cineolifera* |
| RLP173-2 | Old Northern Rd | Si Gel | *P.* sp. Hawkesbury |
| JRN104-2 | Grampians | Dried | *P.* sp. Grampians |
| GW1890-1 | Tapin Tops | Dried | *P. latifolia* |
| NJS424-2 | Tapin Tops | Dried | *P. latifolia* |

#### Table S8 Basic summary statistics for the main species of the project

These diversity statistics are for the species, not mean population statistics. Within species diversity is low mainly because many the loci included are variable only in one or two species. $N$ = number of individuals, $H_O$ = observed heterozygosity, $H_E$ = expected heterozygosity.

| **Species** | $N$ | $H_O$ | $H_E$ |
|---|---|---|---|
| *P. cineolifera* | 50 | 0.053 | 0.067 |
| *P. lanceolata* | 19 | 0.058 | 0.087 |
| *P. ovalifolia* | 21 | 0.032 | 0.076 |
| *P.* sp. Hawkesbury | 13 | 0.067 | 0.079 |
| *P.* sp. Olney State Forest | 6 | 0.062 | 0.072 |

**conStruct analysis including Figs S7, S8, S9**

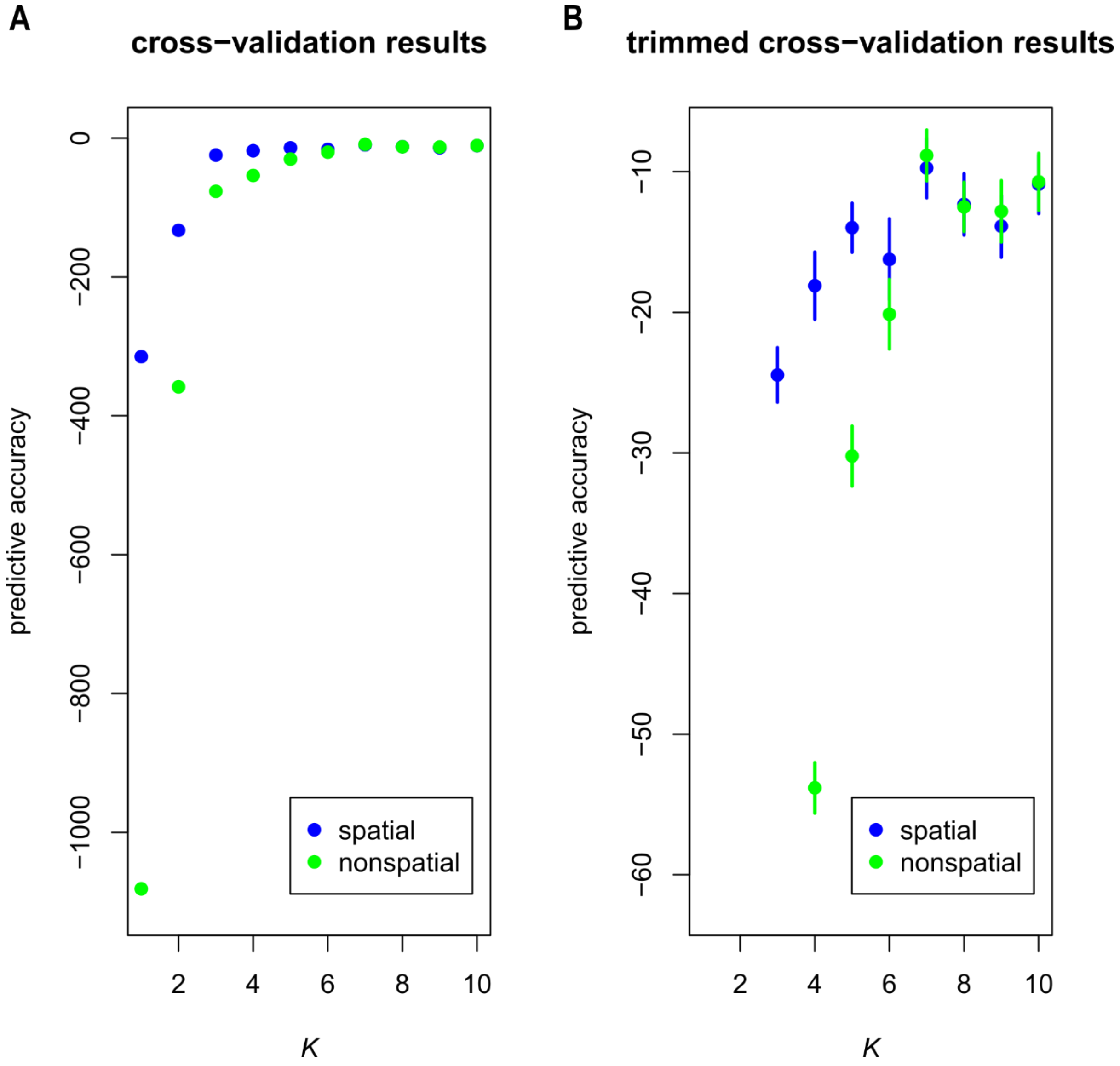


**Fig.S7** ConStruct cross-validation results for the *Prostanthera* final study group[a], comparing spatial and nonspatial conStruct models, run with *K*=1 through 10, with 10 cross-validation replicates. A The results from spatial and nonspatial conStruct models for all values of K. B Zoomed in on the predictive accuracy range from -60 to 0 for the spatial analyses run with *K*=3 through 10.

[a] *P. cineolifera*, *P. lanceolata*, *P. ovalifolia*, *P*. sp. Hawkesbury and *P*. sp. Olney State Forest, TB and PV, *P*. sp. Mt Marsh and P. sp. Oxley Wild Rivers National Park.

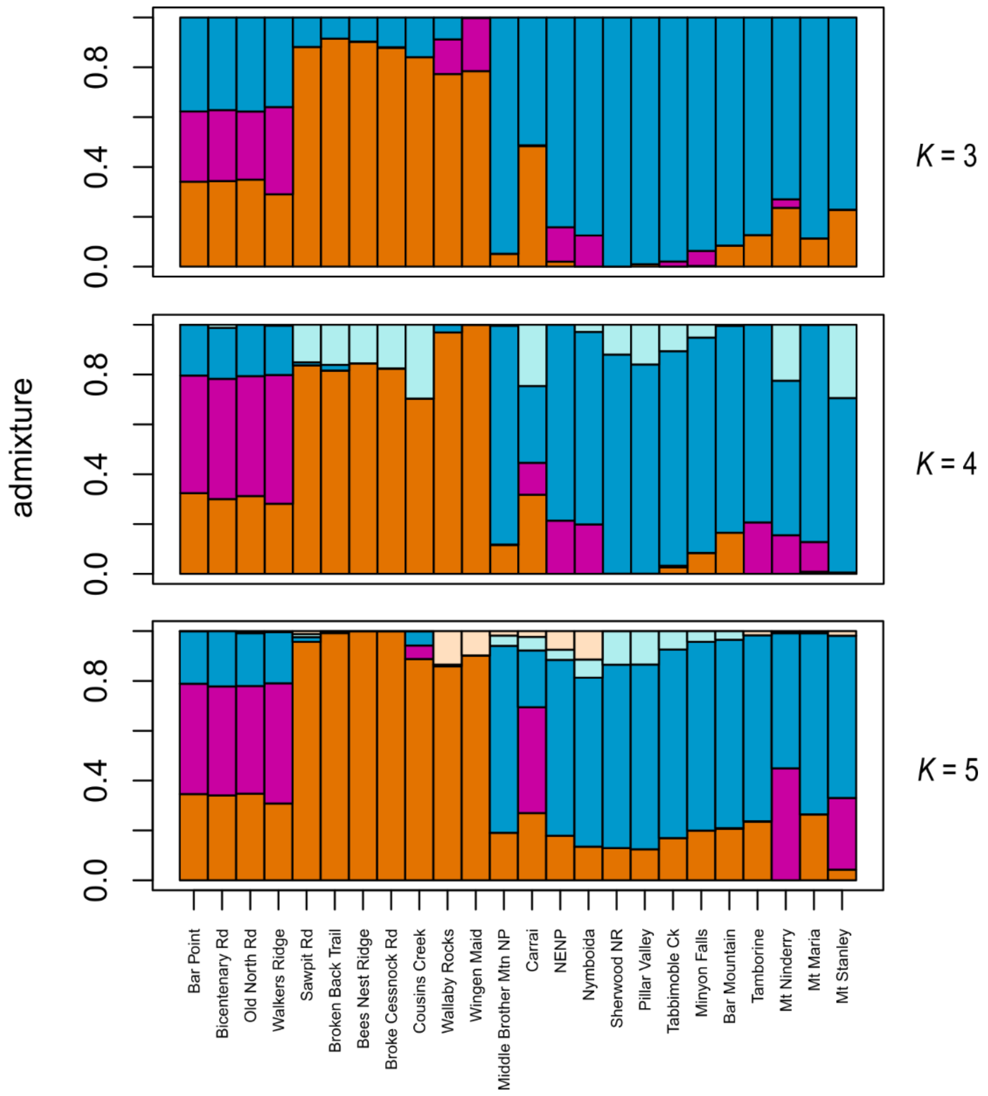


**Fig.S8** Admixture proportions for all populations of *Prostanthera* in the study group estimated using the spatial conStruct model for *K* from 3 to 5.

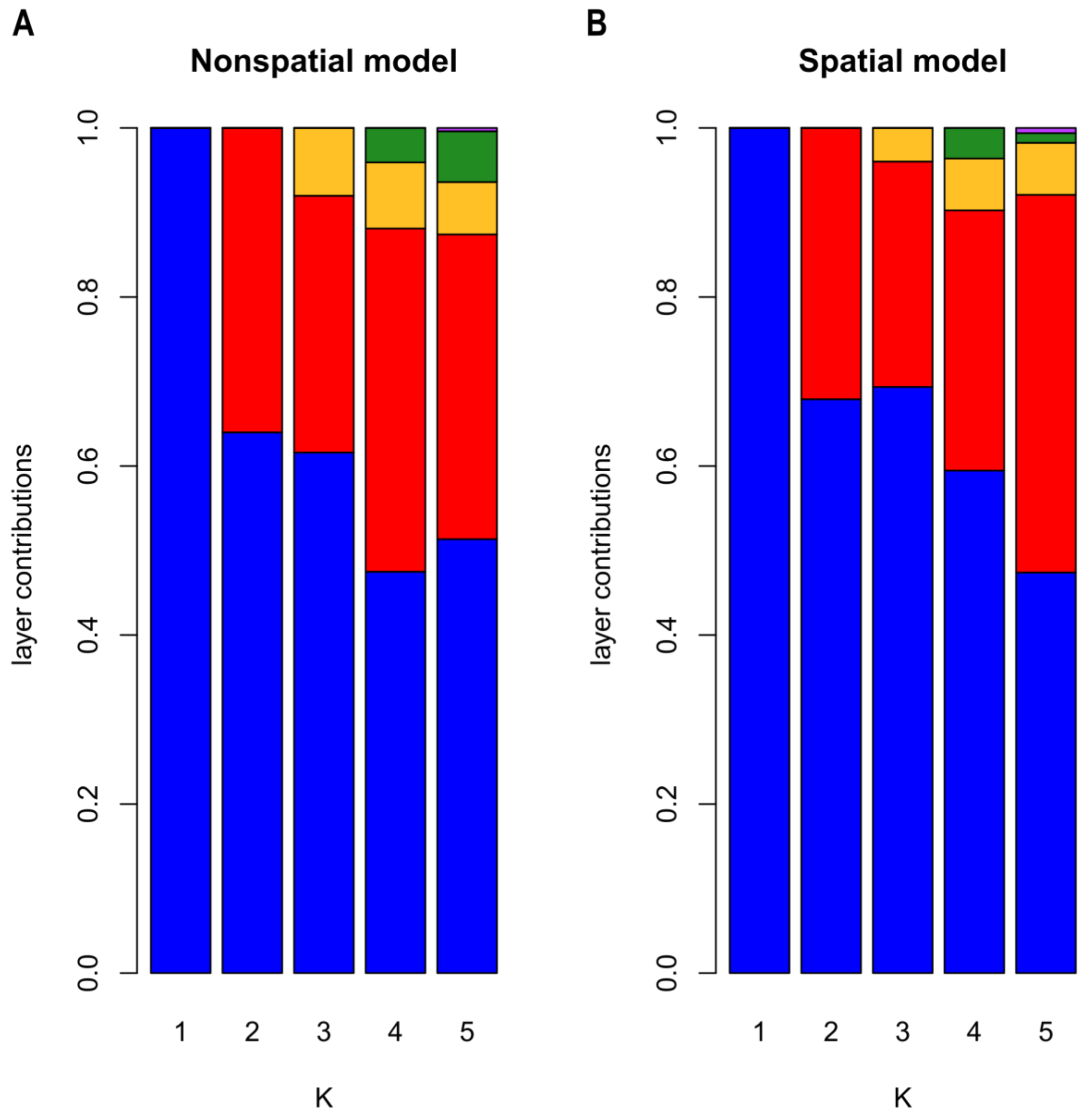


**Fig S9** ConStruct layer/cluster contributions (i.e., how much total covariance is contributed by each layer/cluster), for all layers estimated in runs using K from 1 to 5 for the nonspatial model (A) and the spatial model (B).

#### Extra PCA analyses

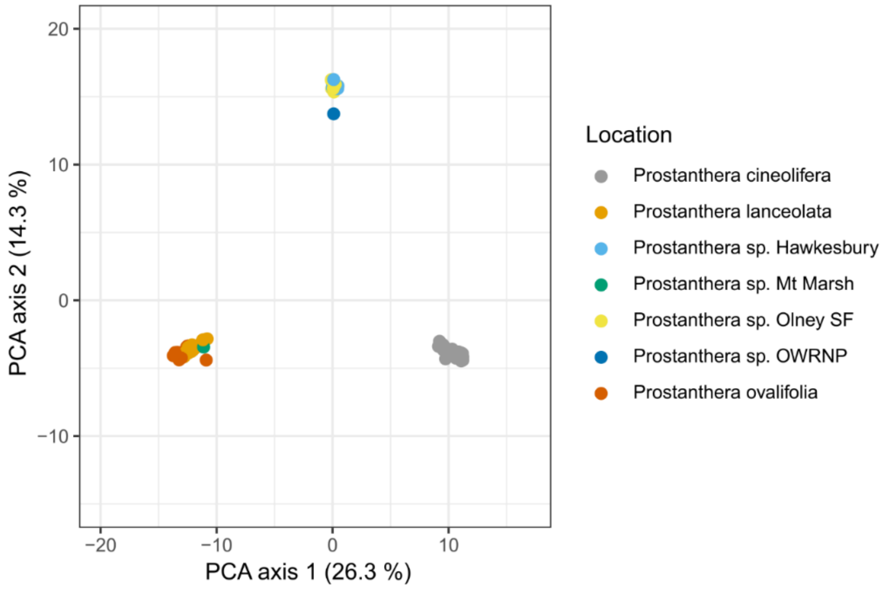


**Fig. S10** Principal Coordinates Analysis (PCA) of SNP data from the main *Prostanthera* dataset (9,559 loci across 122 individuals). Axis 2 vs axis 1. Percentage of genetic distance explained by each of the PCA axes is provided in parentheses. In this analysis, the Tabbimoble Creek and Pillar Valley specimens were included in *P. lanceolata*. *P*. sp. Olney SF = *P*. sp. Olney State Forest, *P*. sp. OWRNP = *P*. sp. Oxley Wild Rivers National Park.

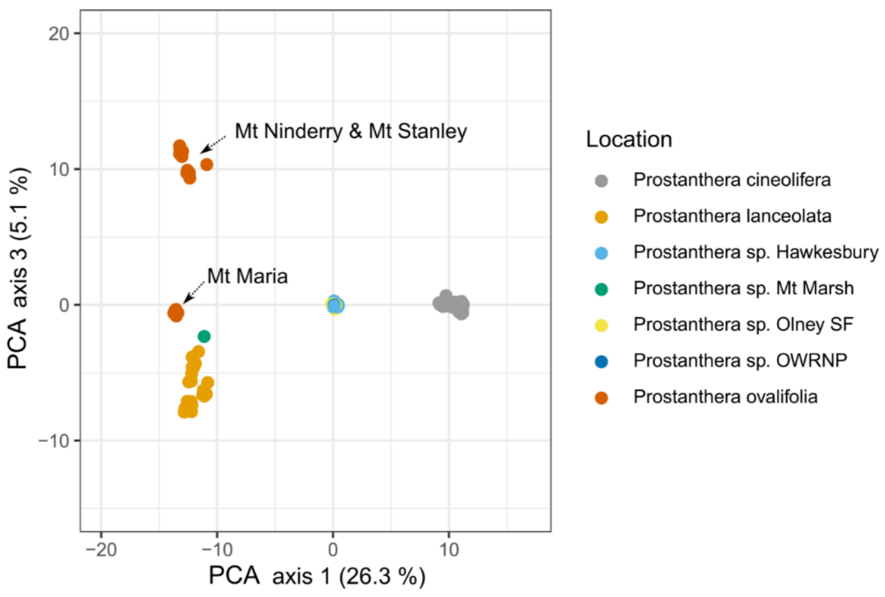


**Fig. S11** Principal Coordinates Analysis (PCA) of SNP data from the main *Prostanthera* dataset (9,559 loci across 122 individuals). Axis 3 vs axis 1. Percentage of genetic distance explained by each of the PCA axes is provided in parentheses. Note: *P*. sp. Oxley Wild Rivers National Park has plotted under *P*. sp. Hawkesbury. In this analysis, the Tabbimoble Creek and Pillar Valley specimens were included in *P. lanceolata*. *P*. sp.

Olney SF = *P*. sp. Olney State Forest, *P*. sp. OWRNP = *P*. sp. Oxley Wild Rivers National Park.

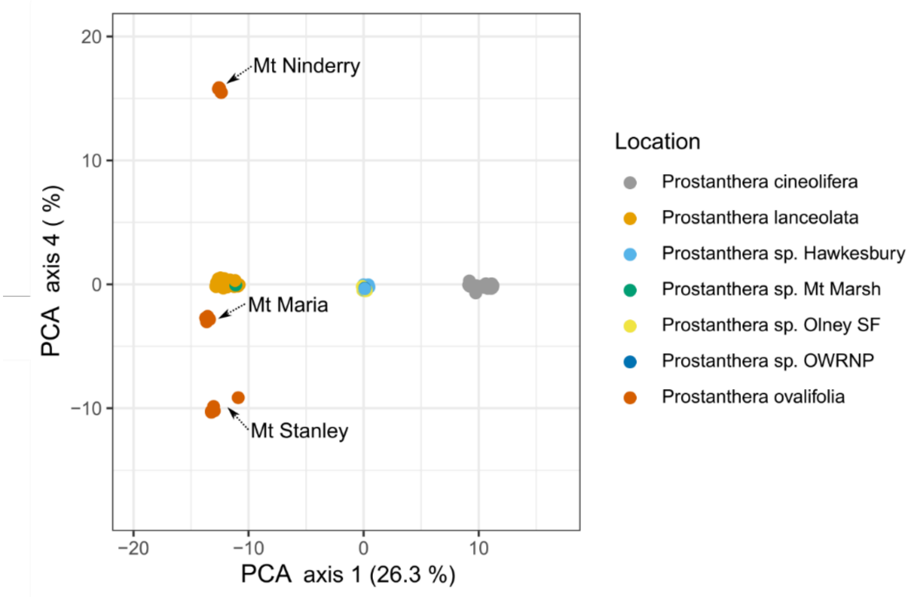


**Fig. S12** Principal Coordinates Analysis (PCA) of SNP data from the main *Prostanthera* dataset (9,559 loci across 122 individuals). Axis 4 vs axis 1. Percentage of genetic distance explained by each of the PCA axes is provided in parentheses. Note: *P*. sp. Oxley Wild Rivers National Park has plotted under *P*. sp. Hawkesbury. In this analysis, the Tabbimoble Creek and Pillar Valley specimens were included in *P. lanceolata*. *P*. sp. Olney SF = *P*. sp. Olney State Forest, *P*. sp. OWRNP = *P*. sp. Oxley Wild Rivers National Park.

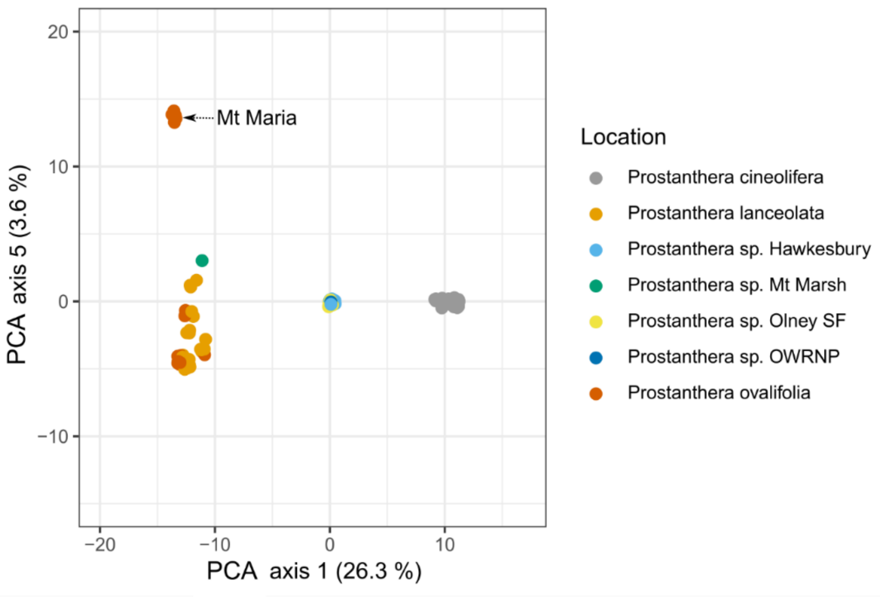


**Fig. S13** Principal Coordinates Analysis (PCA) of SNP data from the main *Prostanthera* dataset (9,559 loci across 122 individuals). Axis 5 vs axis 1. Percentage of genetic distance explained by each of the PCA axes is provided in parentheses. Note: *P*. sp. Oxley Wild Rivers National Park has plotted under *P*. sp. Hawkesbury. In this analysis, the

Tabbimoble Creek and Pillar Valley specimens were included in *P. lanceolata*. *P*. sp. Olney SF = *P*. sp. Olney State Forest, *P*. sp. OWRNP = *P*. sp. Oxley Wild Rivers National Park.

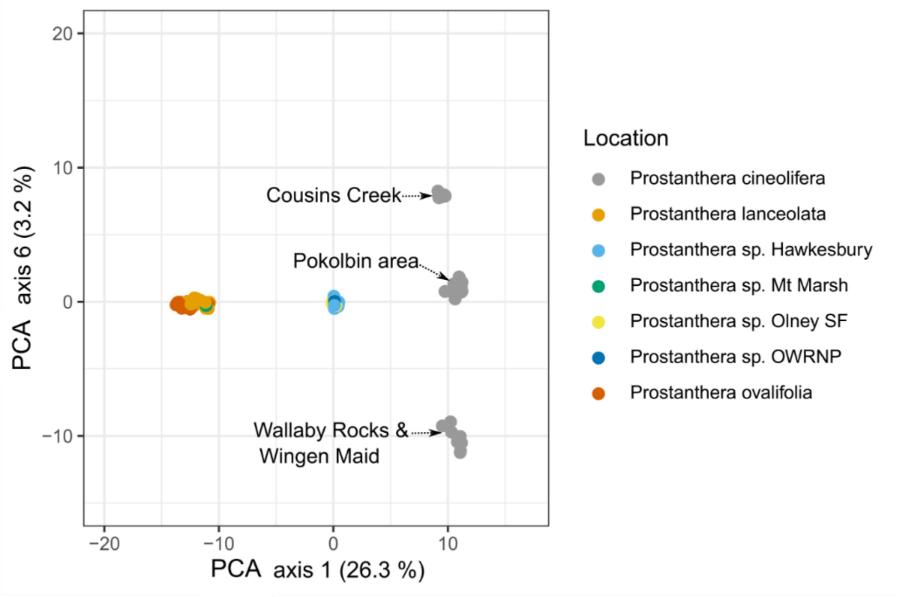


**Fig. S14** Principal Coordinates Analysis (PCA) of SNP data from the main *Prostanthera* dataset (9,559 loci across 122 individuals). Axis 6 vs axis 1. Percentage of genetic distance explained by each of the PCA axes is provided in parentheses. In this analysis, the Tabbimoble Creek and Pillar Valley specimens were included in *P. lanceolata*. *P*. sp. Olney SF = *P*. sp. Olney State Forest, *P*. sp. OWRNP = *P*. sp. Oxley Wild Rivers National Park.

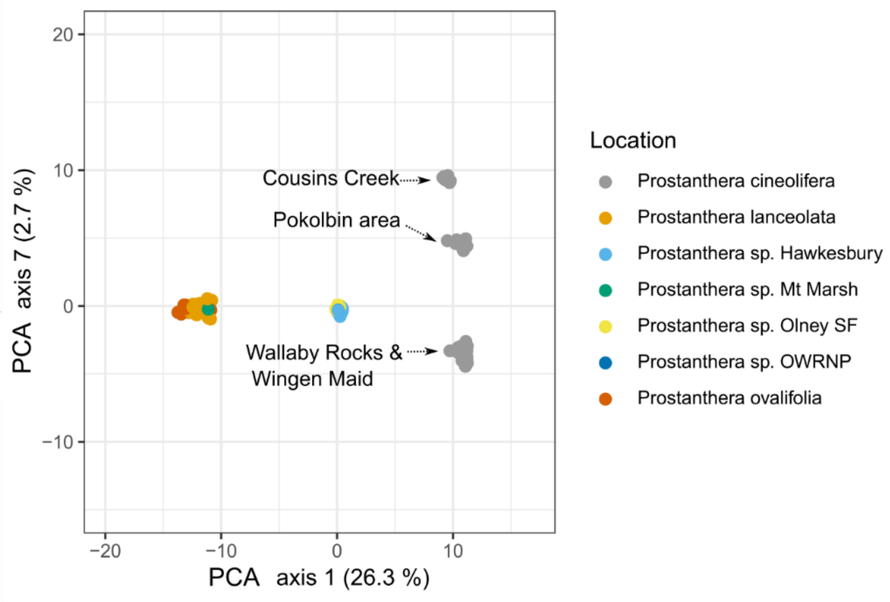


**Fig. S15** Principal Coordinates Analysis (PCA) of SNP data from the main *Prostanthera* dataset (9,559 loci across 122 individuals). Axis 7 vs axis 1. Percentage of genetic distance explained by each of the PCA axes is provided in parentheses. Note: *P*. sp. Oxley Wild Rivers National Park has plotted under *P*. sp. Hawkesbury. In this analysis, the Tabbimoble Creek and Pillar Valley specimens were included in *P. lanceolata*. *P*. sp. Olney SF = *P*. sp. Olney State Forest, *P*. sp. OWRNP = *P*. sp. Oxley Wild Rivers National Park.

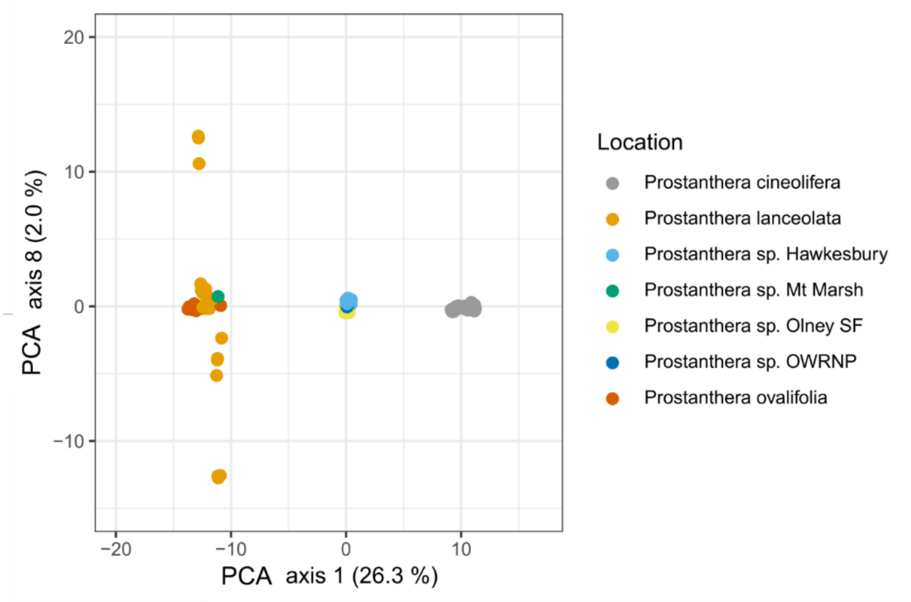


**Fig. S16** Principal Coordinates Analysis (PCA) of SNP data from the main *Prostanthera* dataset (9,559 loci across 122 individuals). Axis 8 vs axis 1. PCA 8 has no effect on samples of *P. cineolifera*, *P. ovalifolia* or the *P.* sp. Hawkesbury group but widely separates samples from four of the eight *P. lanceolata* collection sites. Percentage of genetic distance explained by each of the PCA axes is provided in parentheses. In this analysis, the Tabbimoble Creek and Pillar Valley specimens were included in *P. lanceolata.* *P*. sp. Olney SF = *P*. sp. Olney State Forest, *P*. sp. OWRNP = *P*. sp. Oxley Wild Rivers National Park.

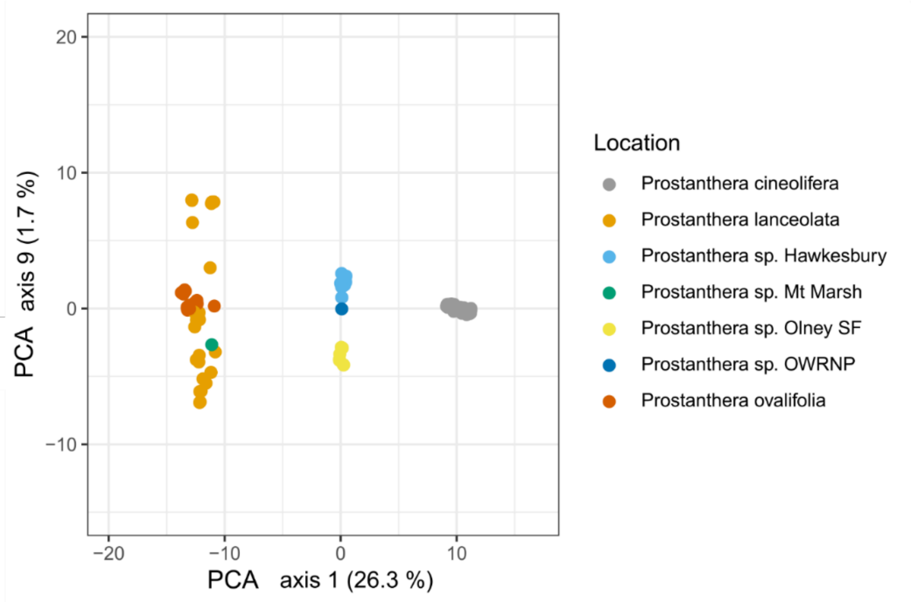


**Fig. S17** Principal Coordinates Analysis (PCA) of SNP data from the main *Prostanthera* dataset (9,559 loci across 122 individuals). Axis 9 vs axis 1. Axis 9 is the first axis to separate samples of the component species of the *P*. sp. Hawkesbury group into separate clusters. Percentage of genetic distance explained by each of the PCA axes is provided in parentheses. In this analysis, the Tabbimoble Creek and Pillar Valley specimens were included in *P. lanceolata*. *P*. sp. Olney SF = *P*. sp. Olney State Forest, *P*. sp. OWRNP = *P*. sp. Oxley Wild Rivers National Park.

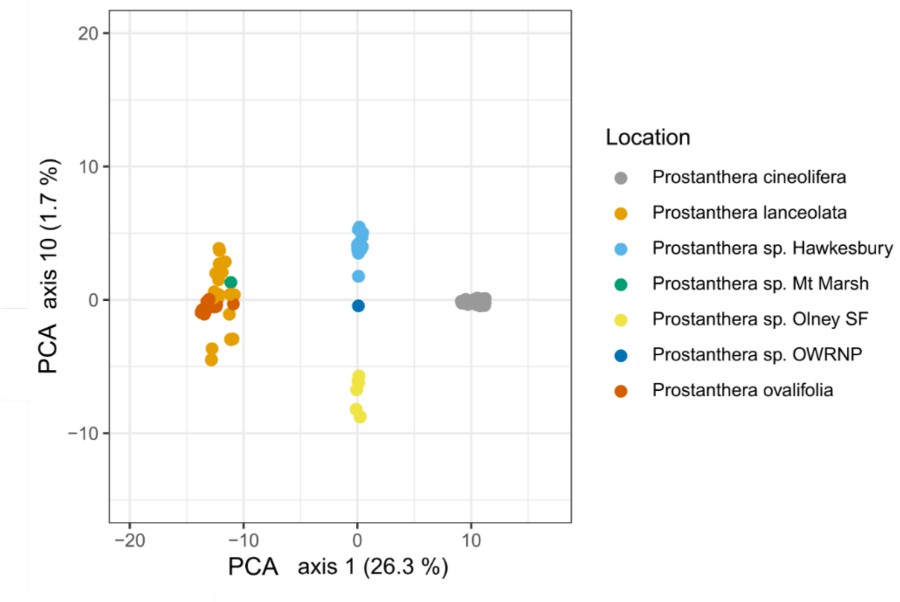


**Fig. S18** Principal Coordinates Analysis (PCA) of SNP data from the main *Prostanthera* dataset (9,559 loci across 122 individuals). Axis 10 vs axis 1. Percentage of genetic distance explained by each of the PCA axes is provided in parentheses. In this analysis, the Tabbimoble Creek and Pillar Valley specimens were included in *P. lanceolata*. *P*. sp. Olney SF = *P*. sp. Olney State Forest, *P*. sp. OWRNP = *P*. sp. Oxley Wild Rivers National Park.

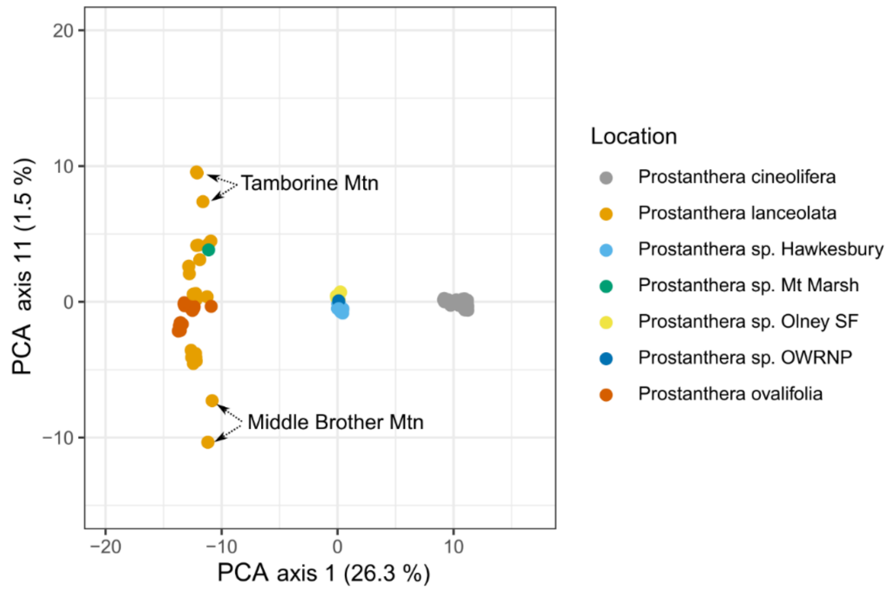


**Fig. S19** Principal Coordinates Analysis (PCA) of SNP data from the main *Prostanthera* dataset (9,559 loci across 122 individuals). Axis 11 vs axis 1. Percentage of genetic distance explained by each of the PCA axes is provided in parentheses. In this analysis, the Tabbimoble Creek and Pillar Valley specimens were included in *P. lanceolata*. *P*. sp. Olney SF = *P*. sp. Olney State Forest, *P*. sp. OWRNP = *P*. sp. Oxley Wild Rivers National Park.

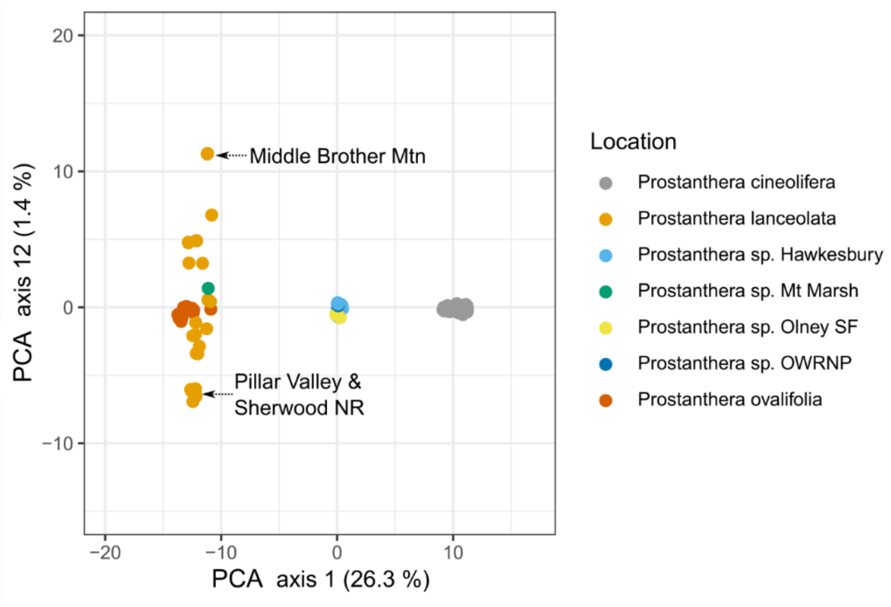


**Fig. S20** Principal Coordinates Analysis (PCA) of SNP data from the main *Prostanthera* dataset (9,559 loci across 122 individuals). Axis 12 vs axis 1. Percentage of genetic distance explained by each of the PCA axes is provided in parentheses. Note: *P*. sp. Oxley Wild Rivers National Park has plotted under *P*. sp. Hawkesbury. In this analysis, the Tabbimoble Creek and Pillar Valley specimens were included in *P. lanceolata*. *P*. sp. Olney SF = *P*. sp. Olney State Forest, *P*. sp. OWRNP = *P*. sp. Oxley Wild Rivers National Park.

**Figs S21, S22 LEA plots**

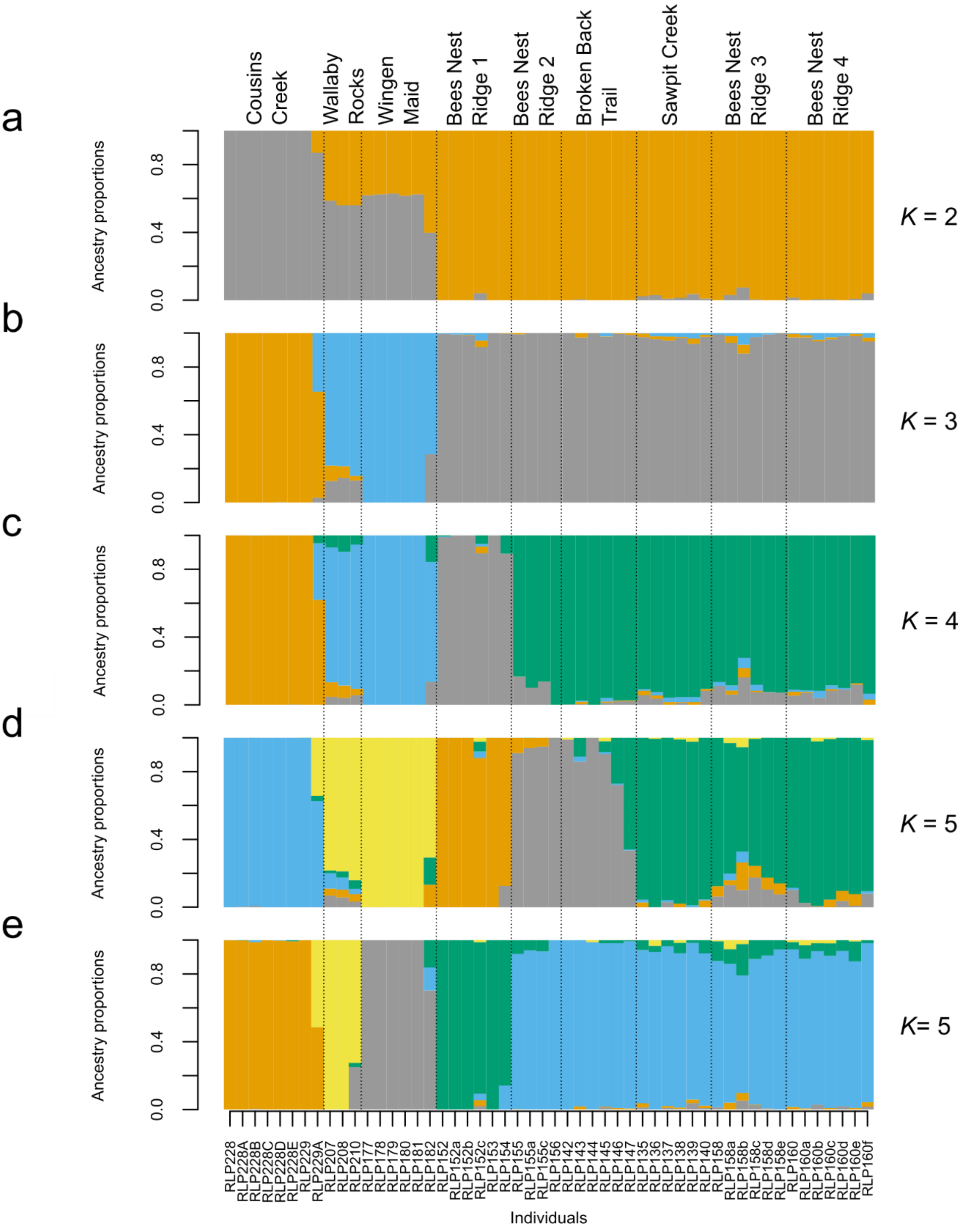


**Fig. S21** Individual ancestry proportions from model-based clustering with sNMF of the *Prostanthera cineolifera* individuals, assuming there are 2–5 genetic clusters/ancestral populations (a-e: $K$ = 2–5, respectively). Note: Each colour only has meaning in one plot; for example, the grey in a is not related to the grey in b, c, d or e. For $K$ = 2 and 3, 10 of 10 replicates indicated clusters as in b and c respectively. For $K$ = 4, nine of 10 replicates indicated clusters as in c. For $K$ = 5, five of 10 replicates indicated clusters as in d, and five of 10 replicates indicated clusters as in e.

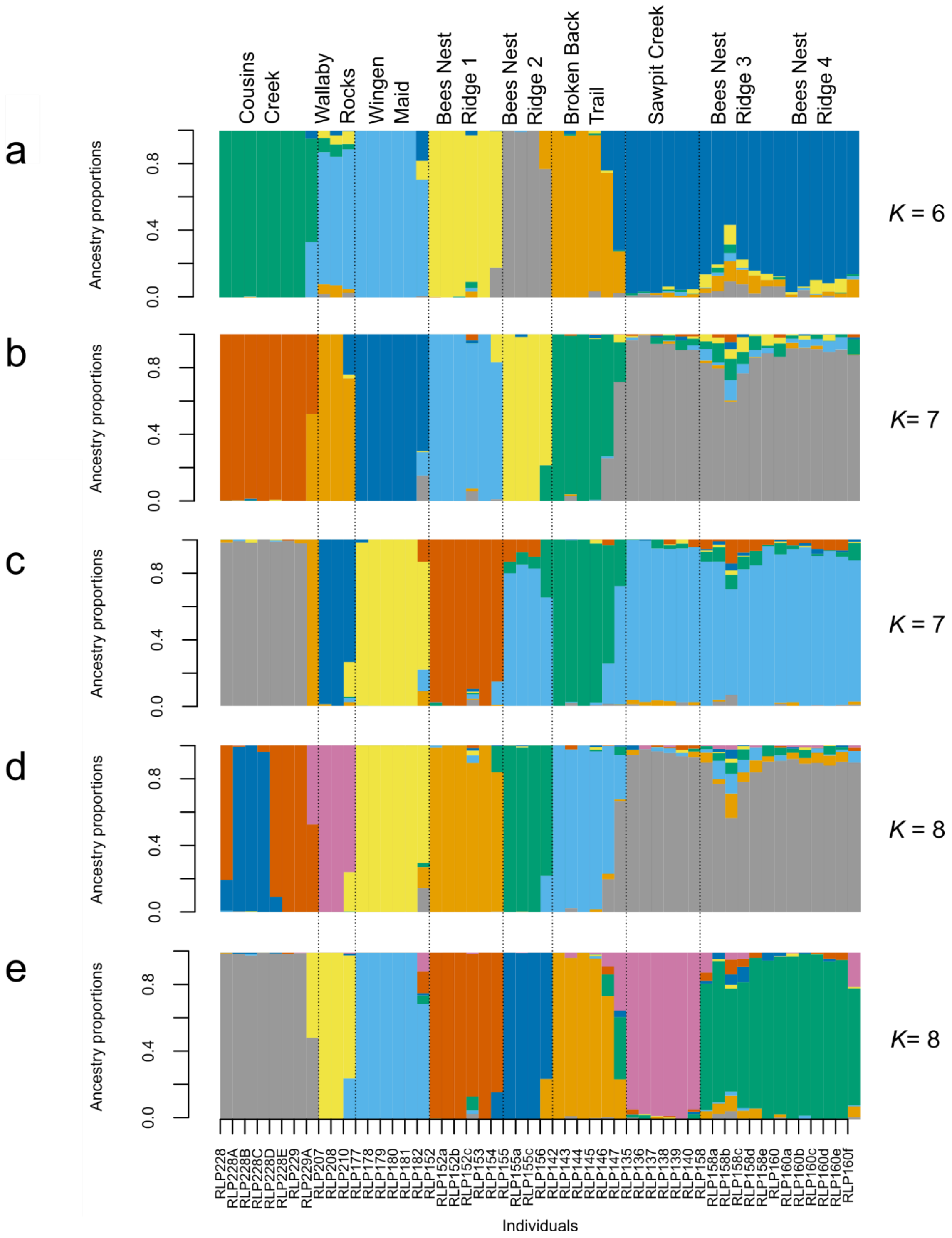


**Fig. S22** Individual ancestry proportions from model-based clustering with sNMF of the *Prostanthera cineolifera* individuals, assuming there are 6–8 genetic clusters/ancestral populations (a-e: $K = 6$–8, respectively). Note: Each colour only has meaning in one plot; for example, the grey in a is not related to the grey in b, c, d or e. For $K = 6$, 10 of 15 replicates indicated clusters as in a. For $K = 7$, four of 10 replicates indicated clusters as in b, and two of 10 replicates indicated clusters as in c. For $K = 8$, three of 10 replicates indicated clusters as in d, and three of 10 replicates indicated clusters as in e.
